# Determination of Slow-binding HDAC Inhibitor Potency and Subclass Selectivity

**DOI:** 10.1101/2021.12.18.473277

**Authors:** Carlos Moreno-Yruela, Christian A. Olsen

## Abstract

Histone deacetylases (HDACs) 1–3 regulate chromatin structure and gene expression. These three enzymes are targets for cancer chemotherapy and are studied for the treatment of immune disorders and neurodegeneration, but there is a lack of selective pharmacological tool compounds to unravel their individual roles. Potent inhibitors of HDACs 1–3 often display slow-binding kinetics, which causes a delay in inhibitor–enzyme equilibration and may affect assay readout. Here, we compare the potency and selectivity of slow-binding inhibitors measured by discontinuous and continuous assays. We find that entinostat, a clinical candidate, inhibits HDACs 1–3 by a two-step, slow-binding mechanism with lower potencies than previously reported. In addition, we show that RGFP966, commercialized as HDAC3-selective probe, is a slow-binding inhibitor with inhibitor constants of 57 nM, 31 nM, and 13 nM against HDACs 1–3, respectively. These data highlight a need for thorough kinetic investigation in the development of selective HDAC probes.

**Table of Contents artwork:** 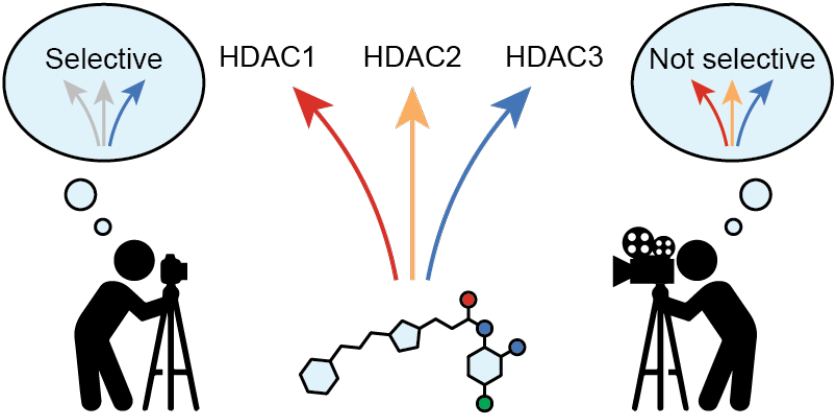

---

Inhibition of enzymes relies on the direct interaction of the inhibitor with its target. Extending the timespan of this interaction, also termed residence time, is pursued in drug design to maximize a compound’s biological impact.^1–2^ Slow-binding inhibitors present low dissociation rates that translate into extended residence times and, often, higher potency.^3–6^ However, slow-binding kinetics also cause a delay in reaching equilibrium within assays, which is not always taken into account in medicinal chemistry campaigns.^1–2^ Disregarding slow-binding mechanisms may lead to an underestimation of affinity against the desired target, but also against off-targets that share structural similarity such as members of the same enzyme family.^1^ Thus, understanding inhibitor kinetics is important for determining potency with accuracy and for assessing subfamily and subclass selectivity.

Histone deacetylases (HDACs) are targeted by chemotherapy, with five HDAC inhibitors in clinical use for the treatment of hematologic cancers.^7^ HDAC inhibitors are also undergoing clinical trials for treatment of dementia and muscular dystrophy^7–8^ and are being investigated in autoimmune diseases.^9^ Humans express 11 Zn^2+^-dependent HDACs, divided into class I (HDACs 1–3 and 8), class IIa (HDACs 4, 5, 7, and 9), class IIb (HDACs 6 and 10), and class IV (HDAC11).^10^ Class I HDACs, the targets of most therapies, remove *N*^ε^-acyllysine posttranslational modifications from nuclear proteins, including histones and transcription factors,^10–14^ and thereby regulate gene expression.^9^ It is often unclear which isozymes are the relevant targets in each disease context, because there is a lack of selective tool compounds to discriminate between class I HDACs. Most inhibitors used as probes inhibit multiple classes (*e.g.* SAHA, **1**, **Figure 1**), are class I-selective (*e.g.* romidepsin, **2**), or inhibit HDACs 1–3 (*e.g.* entinostat, **3**).^15^ Albeit, romidepsin (**2**) is the only recommended probe to study class I HDACs.^16^ Multiple HDAC inhibitors exhibit slow-binding kinetics, first reported for *ortho*-aminoanilides,^17^ but also found for trifluoromethylketones,^18^ hydroxamic acids,^6, 19^ acylhydrazides,^20–21^ as well as romidepsin (**2**).^19, 22^ However, kinetic data is not available for all inhibitors used as probes, such as the claimed HDAC3-selective *ortho*-aminoanilide RGFP966 (**4**).^23^

**Figure 1.**
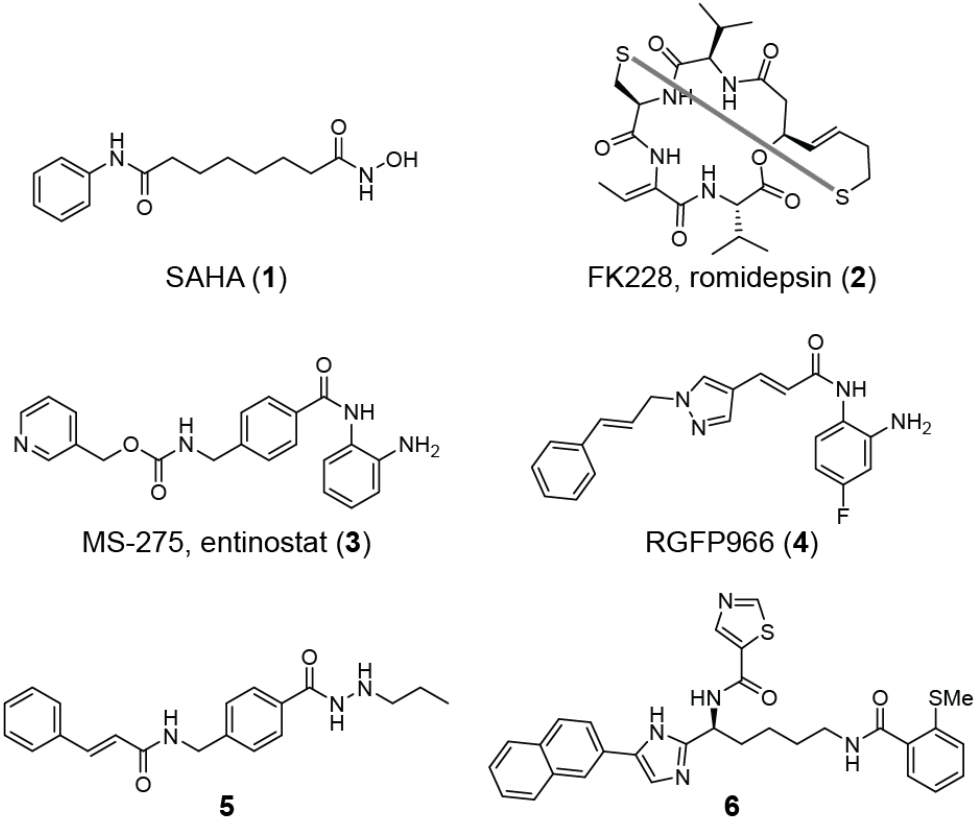
Structures of selected HDAC inhibitors.

Here, we determine the potency and selectivity of slow-binding inhibitors of HDACs 1–3 using standard end-point assays as well as discontinuous and continuous experiments that account for slow inhibitor–enzyme equilibration. We find that discontinuous assays often underestimate potency and that the calculated selectivity is highly dependent on the assay format. Moreover, our data show that RGFP966 (**4**) is not an HDAC3-selective probe.

We selected four inhibitors for our investigations; SAHA (**1**), entinostat (**3**), RGFP966 (**4**), and acylhydrazide **5**.^24^ Compounds **1** and **3** inhibit HDACs 1–3 with similar potency in end-point assays, while they display different kinetics of inhibition (fast- and slow-binding kinetics, respectively).^15, 17, 25^ Compounds **4** and **5**, on the other hand, are reported to exhibit selectivity for HDAC3^23–24^ but their binding kinetics have not been investigated although they are structurally similar to known slow-binding inhibitors.^7, 21^

First, we evaluated inhibition of HDACs 1–3 using a standard fluorescence assay based on a 7-amino-4-methyl-coumarin (AMC)-coupled fluorogenic substrate.^15, 26^ The substrate, Ac-Leu-Gly-Lys(Ac)-AMC, has similar Michaelis-Menten constants (*K*_M_) for each enzyme in the low micromolar range, which allows investigation at comparable enzyme saturation conditions.^15^ Inhibitor and substrate were incubated with the enzyme for 30 min before development with a protease, providing end-point dose–response curves (**Figure 2A**). SAHA (**1**) and entinostat (**3**) showed similar potency against the three enzymes, while RGFP966 (**4**) and compound **5** showed selectivity for HDAC3 as previously reported. However, the measured potency and selectivity of RGFP966 (**4**) differed significantly from the literature, with an IC_50_ of 514 ± 4 nM against HDAC3 (compared to 80 nM) and a selectivity ratio of 3.5 vs. HDAC2 (compared to >180) (**Figure 2D**).^23^ Data for compound **5** was in better agreement with previous reports, with a 6.1 selectivity ratio vs. HDAC1 (compared to 12.4).^24^

**Figure 2.**
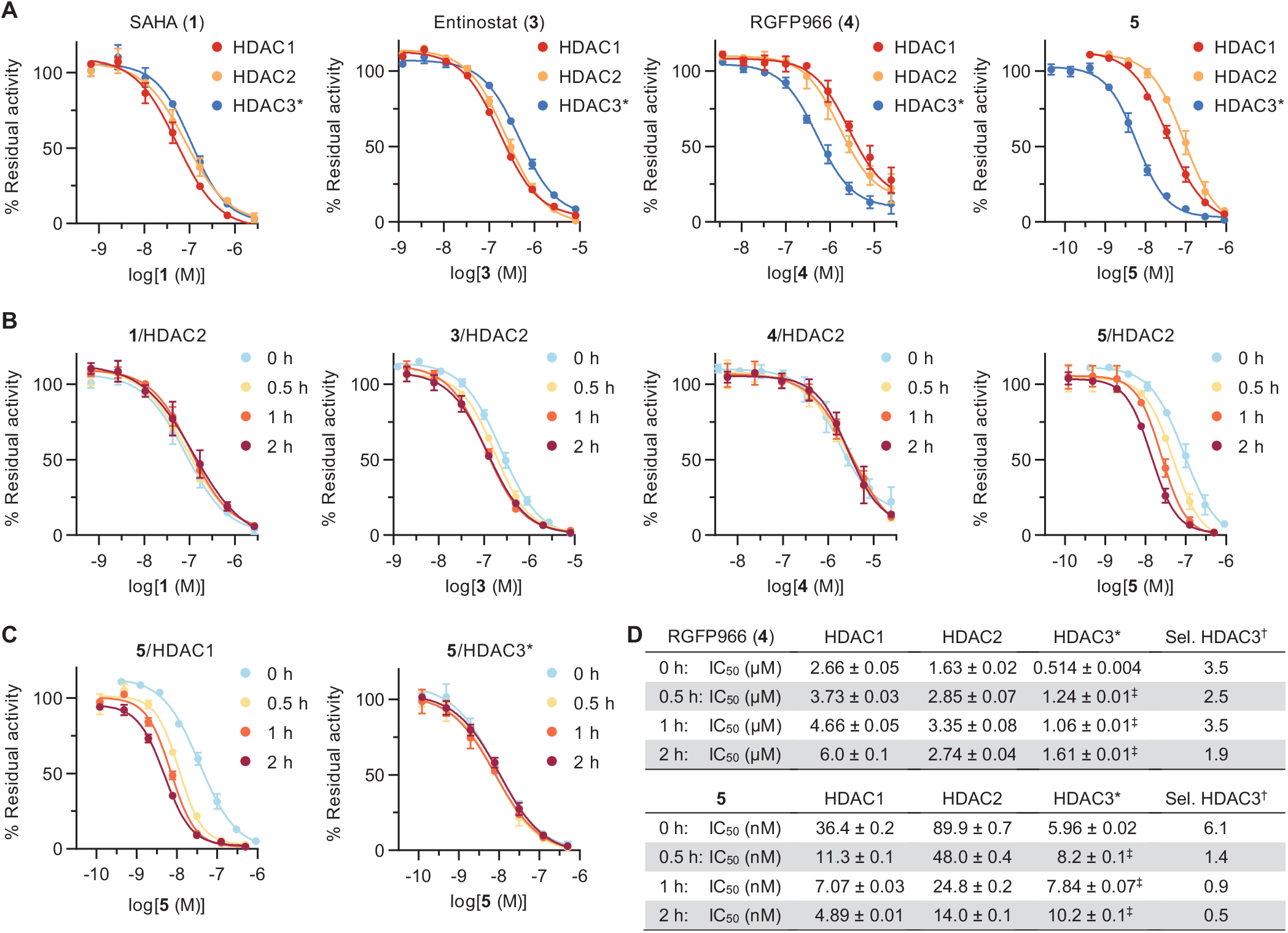
Discontinuous inhibition of HDACs 1–3. (A) Dose-response curves measured after 30 min reaction with enzyme and substrate. (B) HDAC2 curves after pre-incubation with enzyme for 0, 0.5, 1, or 2 h, followed by addition of substrate and 30 min reaction. (C) HDAC3 curves of compounds **3** and **5** with and without pre-incubation, measured in buffer without reducing agents or surfactants. See **Supporting Figure S1A,B** for HDAC1 and additional HDAC3 curves. (D) IC50 data of compounds **4** and **5**, and selectivity of HDAC3 inhibition. See **Supporting Figure S1C** for data for compounds **1** and **3**. All data represent mean ±SD, *n* ≥ 2. *HDAC3 incubated with the deacetylase activation domain (DAD) of NCoR2. ^†^Selectivity calculated vs. the second most inhibited enzyme (**4**: HDAC2, **5**: HDAC1), by transforming IC_50_ data into *K*_i_ and calculating the ratio (see data analysis in the Supporting Information). ^‡^Data obtained in buffer without reducing agents or surfactants (see **Supporting Figure S2** and pre-incubation methods for choice of buffer).

Compounds **3**, **4**, and **5** contain either *ortho*-aminoanilide or acylhydrazide Zn^2+^-binding groups, which are prevalent in inhibitors of class I HDACs that exhibit slow-binding kinetics.^17, 20, 25^ Since slow-binding inhibitors commonly equilibrate after periods of time longer than 30 min,^27^ the performed end-point assays would not be expected to occur at steady-state.

To enable equilibration, assays were next performed with pre-incubation of inhibitor and enzyme during 30 min, 1 h or 2 h before substrate addition. SAHA (**1**), an inhibitor that exhibits fast-on/fast-off binding kinetics, inhibited HDACs 1 and 2 with similar potency within experimental error regardless of pre-incubation. The same trend was observed for entinostat (**3**) and RGFP966 (**4**) with only slight changes in IC_50_ values in response to pre-incubation, which may indicate inhibitor–enzyme equilibration at the time scale 0–30 min (**Figure 2B,D** and **Supporting Figure S1**). Only compound (**5**) followed a slow-binding profile for HDACs 1 and 2 inhibition, with substantial decrease in IC_50_ values upon pre-incubation (**Figure 2B–D**).The selectivity of compounds **4** and **5** were calculated based on each pre-incubation dataset. Compound **4** maintained a similar selectivity towards HDAC3 at all pre-incubation times, with just a slight loss in selectivity at 2 h, which is within the experimental error of the system (1.9 selectivity ratio vs. HDAC2, **Figure 2D**). More strikingly, the selectivity of compound **5** dropped from 6.1 to 1.4 after 30 min and further decreased at the 1 h and 2 h time-points, due to a decrease in IC_50_ for HDACs 1 and 2 while potency against HDAC3 was maintained. Since HDAC3 experiments were carried out at 1–5 nM enzyme concentration, this difference in behavior might be explained by compound **5** already approaching stoichiometric inhibition of HDAC3 (tight binding) without pre-incubation. Then, potential further increase in potency after pre-incubation would thus be detected for HDACs 1 and 2 only. Nonetheless, the data highlighted the need for more thorough investigation of slow-binding inhibitor potency to assess selectivity with better accuracy.

Pre-incubation data of slow-binding inhibitors can be fit to exponential decay functions in order to calculate slow-binding kinetic and inhibitor constants.^27^ Here, this analysis could be performed for entinostat (**3**) against HDACs 2 and 3, and for compound **5** against HDACs 1 and 2, providing a rough estimate of compound potency and residence time (**Supporting Figure S3**). Through this analysis, entinostat (**3**) was calculated to have apparent inhibition constants of ~73 nM (HDAC2) and ~27 nM (HDAC3), while compound **5** exhibited even lower potency. Unfortunately, selectivity could not be recalculated due to the lack of complete data sets for HDACs 1–3.

Next, we adapted the initial discontinuous assay conditions to a continuous format, by adding the protease developer directly to the reaction as previously reported.^17, 28^ Protease concentration was optimized to obtain linear substrate conversion by the HDAC for 40–60 min, while ensuring deacetylation by the HDAC was the rate-limiting step (**Supporting Figure S4**).^29^ With these assay conditions in hand, we measured enzyme kinetics to determine the Michaelis-Menten constant for each of our enzyme preparations (**Supporting Figure S5**), and recorded enzyme activity at different concentrations of inhibitor. Fast-on/fast-off inhibitors such as SAHA (**1**) equilibrate rapidly to steady-state to afford linear progression curves at all inhibitor concentrations.^6, 17–18^ Conversely, slow-binding inhibitors lower the reaction rate over time, resulting in bending assay progression curves. Fitting the apparent first-order kinetic constant of equilibration (*k*_obs_) to a linear or a hyperbolic function of inhibitor concentration reveals whether the inhibitor follows mechanism A or mechanism B of slow-binding kinetics, respectively (**Figure 3A**). In mechanism A, a single step of binding of the inhibitor to the enzyme is detected, with slow overall on- and off-rates. In the more common mechanism B, rapid formation of an initial inhibitor–enzyme complex is found, followed by a slow transition to a more stable and long-lived enzyme–inhibitor complex (EI*).^27, 30^

**Figure 3.**
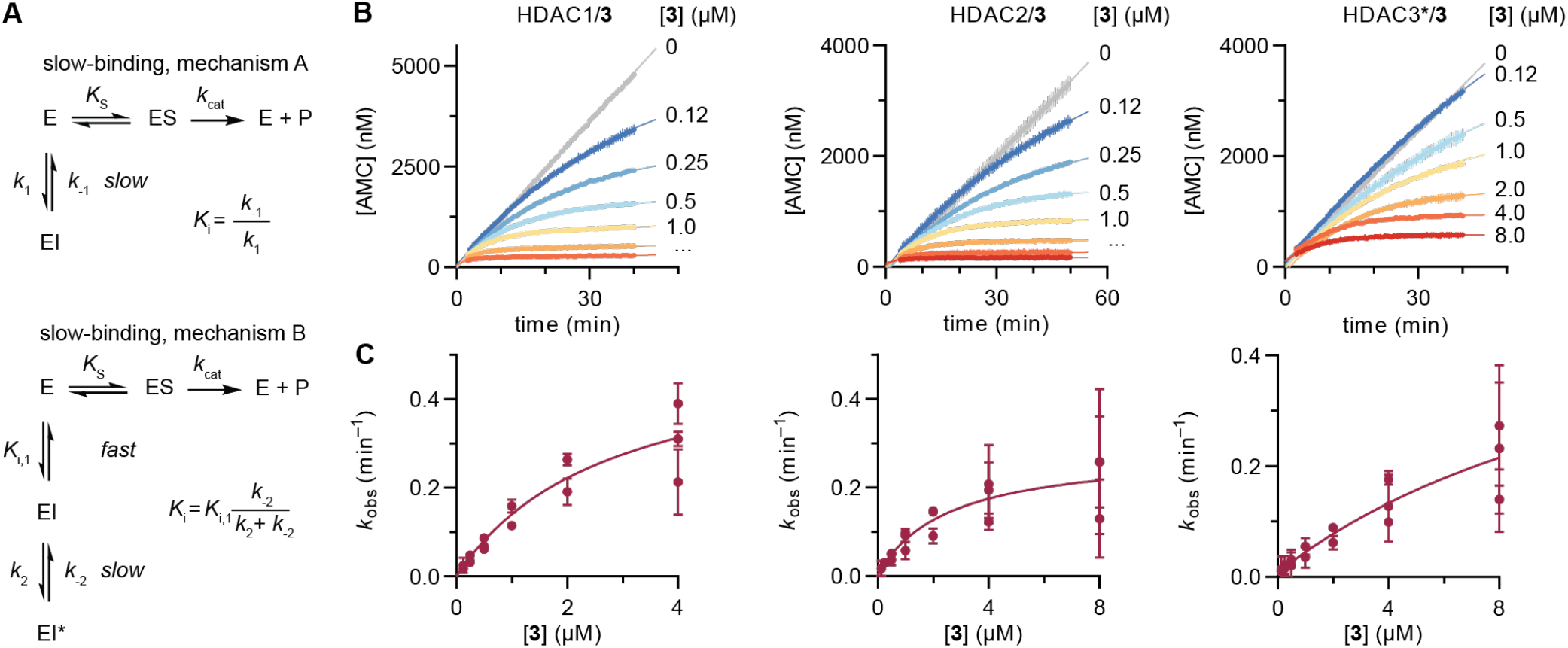
Models of slow-binding inhibition and continuous inhibition of HDACs 1–3 by entinostat (**3**). (A) Mechanism A and mechanism B of competitive slow-binding inhibition, and calculation of inhibitor constants (*K*_i_) from kinetic data. (B) Continuous assay progression curves for the inhibition of HDACs 1–3 by different concentrations of compound **3**. (C) Secondary plots of the apparent first-order rate constant of equilibration (*k*_obs_) vs. inhibitor concentration, and fitting to mechanism B of slow kinetics (see **Table 1** for numerical data). Data represent mean ±SEM of individual experiments, with each experiment performed at least twice. *The HDAC3 preparation contains the DAD of NCoR2.

Entinostat (**3**) afforded the characteristic bending progression curves of a slow-binding inhibitor against all three HDACs (**Figure 3B**). The *k*_obs_ data fitted well to mechanism B of HDAC inhibition (**Figure 3C**, hyperbolic relationship), which adds further insight to the previous kinetic analysis of this compound.^25, 31^ Data revealed that entinostat (**3**) presents a fast first binding step with equilibrium constants (*K*_i,1_) of ~0.59 μM for HDACs 1 and 2, and ~3.2 μM for HDAC3, which are similar to previous estimations of potency.^25, 31^ Taking all kinetic constants into account, the calculated potency (*K*_i_) of entinostat (**3**) was <1 nM against HDAC1, ~6 nM against HDAC2, and ~39 nM against HDAC3 (**Table 1**). These values are much lower than those obtained from end-point experiments by us and others,^32^ including HDAC2 pre-incubation assays, and also than those calculated by previous fitting to mechanism A of slow binding.^25, 31^ Our kinetic data may help explain the high potency of entinostat (**3**) in cellular assays, as well as its long-lasting effect due to extended enzyme dissociation half-life (t_½_, **Table 1**).^25, 32^

**Table 1.**
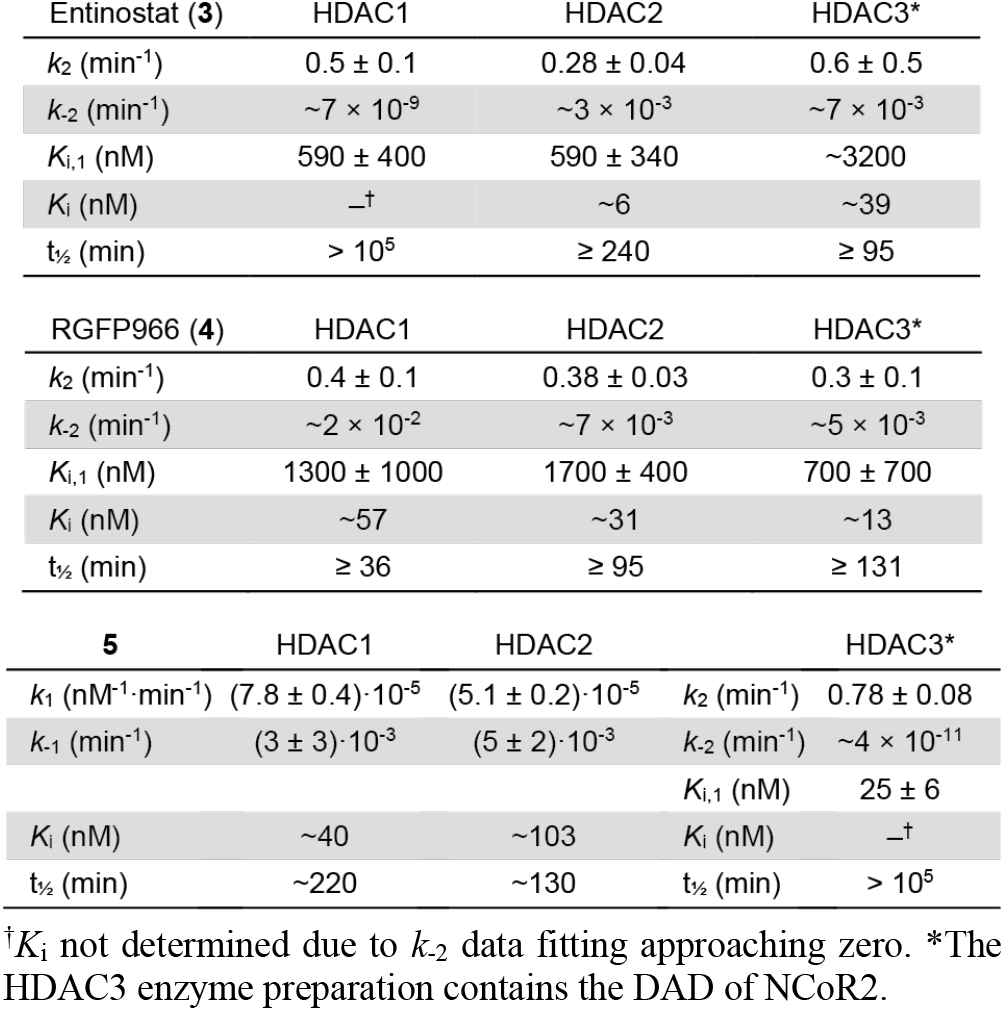
Calculated kinetic constants (*k*_n_), inhibitor constants (*K*_n_), and dissociation half-lives (t_½_) of compounds **3**, **4**, and **5**. Data corresponds to mechanism B of slow-binding kinetics, except for the inhibition of HDACs 1 and 2 by compound **5**, which follows mechanism A.

Continuous assays with RGFP966 (**4**) revealed slow inhibition of HDACs 1–3, which was anticipated based on its chemical structure but was not indicated in the discontinuous assays discussed above. Bending of assay progression curves was especially prominent for HDAC3 (**Figure 4A**), which emphasizes the higher sensitivity of continuous assays towards detecting slow-binding profiles. Inhibition of HDACs 1–3 followed mechanism B of slow-binding kinetics and afforded inhibitor constants of ~57 nM for HDAC1, ~31 nM for HDAC2, and ~13 nM for HDAC3 (**Table 1**). Based on these results, RGFP966 (**4**) exhibits a mere 2.4-fold selectivity towards HDAC3, which is similar to that obtained from pre-incubation (2 h time-point, **Figure 2D**). On the other hand, the calculated potency against each enzyme is >10 times lower than end-point estimates. Overall, our data indicates that RGFP966 (**4**) is a potent inhibitor of HDACs 1–3 with only a minor preference for HDAC3 inhibition.

**Figure 4.**
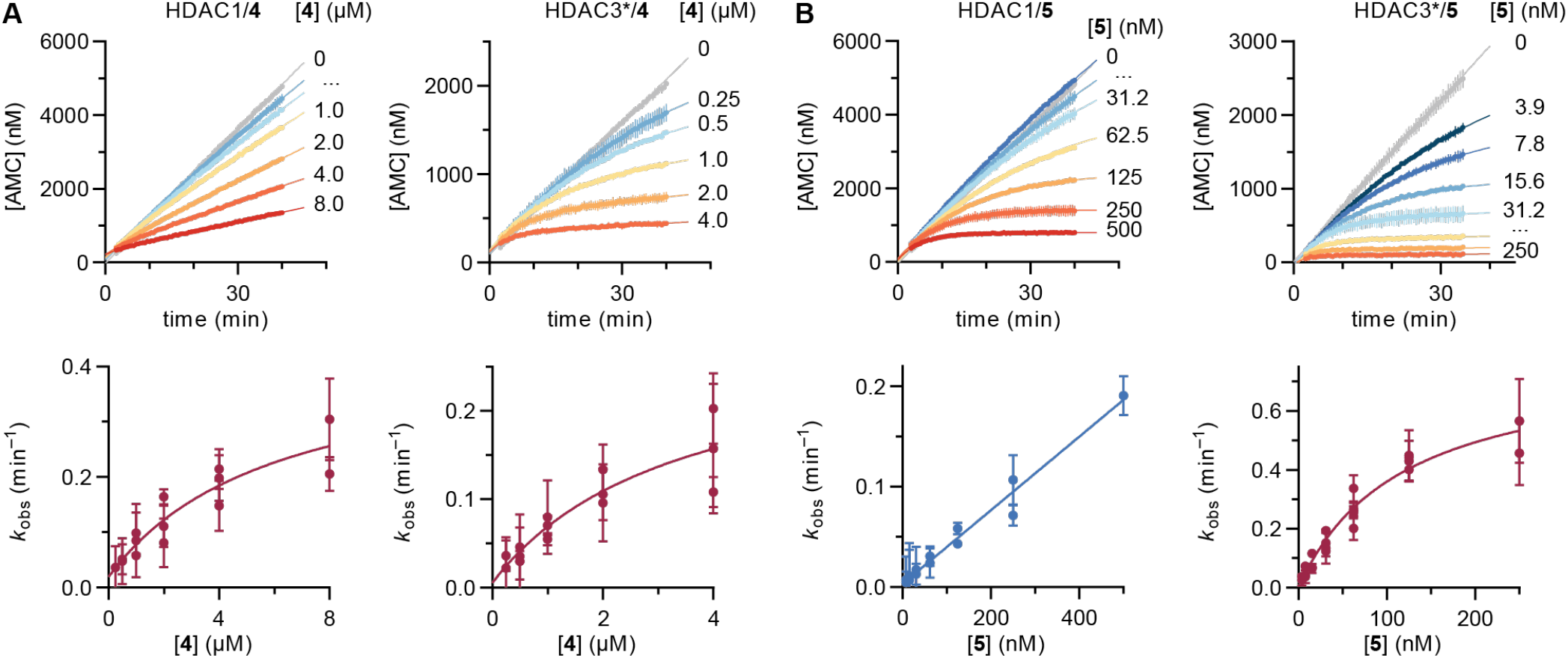
Continuous inhibition of HDACs 1 and 3 by RGFP966 (**4**) and compound **5**. (A) Continuous assay progression curves (top graphs) and *k_obs_* secondary plots (bottom graphs, mechanism B) relative to compound **4**. (B) Continuous assay progression curves (top graphs) and *k_obs_* secondary plots (bottom graphs, HDAC1 data fitted to mechanism A and HDAC3 data fitted to mechanism B of slow binding) relative to compound **5**. Data represent mean ±SEM of individual experiments, with each experiment performed at least twice. See **Supporting Figure S6** for HDAC2 data and **Table 1** for numerical data. *The HDAC3 preparation contains the DAD of NCoR2.

The slow-binding behavior of compound **5** against HDACs 1 and 2, which was identified by pre-incubation, was recapitulated by continuous assays. In addition, we also detected slow-binding inhibition of HDAC3 (**Figure 4B**).

Interestingly, secondary plots of *k*_obs_ indicated that compound **5** follows mechanism A of HDAC1,2 inhibition, whereas it inhibits HDAC3 through mechanism B of slow binding (**Figure 4B** and **Supporting Figure S6B**). These differences in kinetic mechanism are not uncommon and were reported previously for a trifluoromethylketone inhibitor^18^ as well as the natural product trapoxin A.^6^ As a result of data fitting, the calculated *K*_i_ values of compound **5** against HDACs 1 and 2 were ~40 nM and ~103 nM, respectively (**Table 1**), which are somewhat higher than indicated by the IC_50_ values obtained by end-point experiments (**Figure 1D**).^24^ Conversely, the data for inhibition of HDAC3 indicated pseudo-irreversible inhibition with very low off-rates (*k*_-2_ ~0), and a first step *K*_i,1_ of 25 ±6 nM. Thus, compound **5** is highly potent against HDAC1–3, with enzyme residence times of more than 2 h (**Table 1**), and with slow, tight-binding behavior of HDAC3 inhibition. Future studies will reveal whether this kinetic profile could be exploited towards selective inhibition of HDAC3 in a biological setting.

HDACs 1–3 form multiprotein complexes with diverse epigenetic functions^33^ and reports have shown that inhibitor potency and selectivity may differ between the free and the complexed HDAC forms.^34–35^ Thus, inhibitor assays where the HDAC is accompanied by a complex partner likely produces different results of inhibition than free recombinant enzymes. Commercial HDAC3 includes the HDAC-interacting DAD domain of NCoR2, which is required for enzymatic activity, and this interaction can be further stabilized *in vitro* by inositol phosphates.^36^ Since standard experiments are performed with HDAC3 and the DAD in dynamic equilibrium, we also studied the effect of adding inositol hexaphosphate (InsP_6_) to HDAC3 experiments, at a concentration shown to fully stabilize the complex.^37^ End-point inhibition data without pre-incubation remained similar to data recorded without InsP_6_ added. Conversely, pre-incubation of inhibitor with enzyme and InsP_6_ provided stronger inhibition with 10–20 times lower IC_50_ values for entinostat (**3**), RGFP966 (**4**), and compound **5** after 2 h (**Figure 5A** and **Supporting Figure S7A,B**). Addition of InsP_6_ enhanced HDAC activity and the experiments could be performed at 5 times lower enzyme concentration but compound **5** still afforded stoichiometric (tight binding) inhibition. These data suggest that the three compounds inhibit the activity of a stabilized HDAC3/NCoR2 complex with higher potency and/or slower kinetics compared to the more dynamic standard enzyme preparation. Such behavior appears different from the binding studies reported for the *ortho*-aminoanilide Cpd-60, which presents lower affinity for a pre-stabilized HDAC3/NCoR2 complex using InsP_6_.^37^

**Figure 5.**
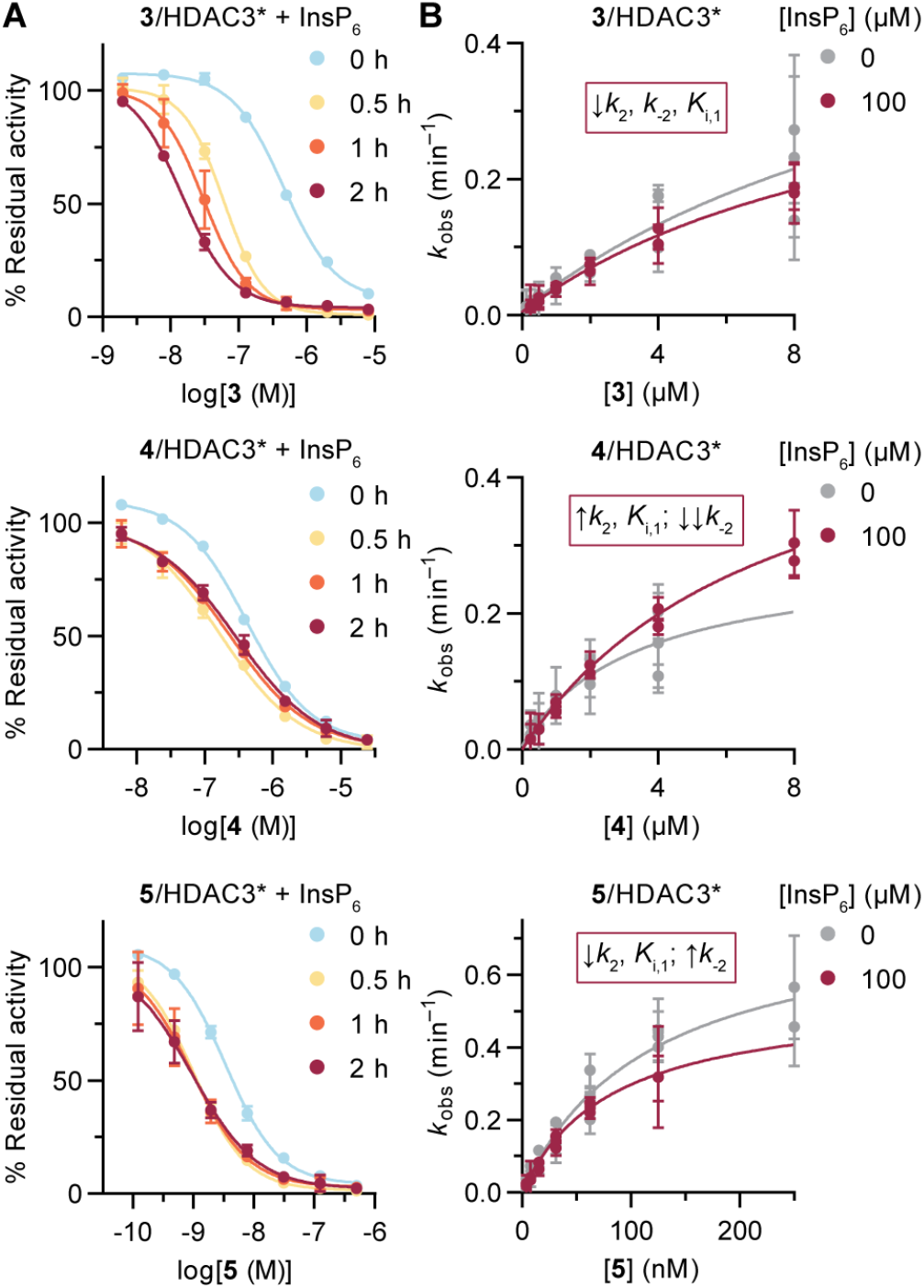
Inhibition of HDAC3 in the presence of inositol hexaphosphate (InsP_6_). (A) Curves of compounds **3**, **4** and **5** with and without pre-incubation, measured in buffer without reducing agents or surfactants. See **Supporting Figure S7A,B** for curves of compound **1** and for IC_50_ and selectivity data. Data represent mean ±SD, *n* = 2. (B) *k*_obs_ secondary plots relative to compounds **3**, **4** and **5** with and without InsP_6_. Data represent mean ± SEM of individual experiments, with each experiment performed at least twice. See **Supporting Figure S7C,D** for progression curves and numerical data. A summary of data changes with InsP_6_ is provided for each compound. *The HDAC3 preparation contains the DAD of NCoR2.

We then performed continuous inhibition assays with compounds **3**, **4**, and **5**, where the HDAC3/DAD preparation was pre-mixed with InsP_6_ before addition to the assay plate. The impact of complex stabilization was less pronounced in these experiments. All three inhibitors maintained mechanism B of slow-binding kinetics; albeit, with changes in the kinetic constants being were observed in each case (**Figure 5** and **Supporting Figure S7C,D**). Entinostat (**3**) gained affinity in the first binding step and showed slower kinetics for the transition to the more stable EI* complex, affording an estimated *K*_i_ of 24 nM. RGFP966 (**4**) exhibited a decrease in affinity for the first step but gained substantial overall potency due to very low off-rates (*k*_-2_ ~0). Lastly, compound **5** gained affinity in the first step and remained a slow, tight-binding inhibitor of the stabilized HDAC3/DAD complex.

Our data underpins the need for detailed kinetic characterization of potent class I HDAC inhibitors, with particular importance when determining subclass selectivity. RGFP966 (**4**) is reported and commercialized as HDAC3-selective probe and has been employed in numerous studies currently associated with specific HDAC3 biology.^23, 38–40^ Here, we show that this compound is not appropriate as an HDAC3-selective probe, which calls into question the biological functions assigned to HDAC3 based on the use of RGFP966 (**4**).

A recent report by Liu and coworkers identified compound **6** (**Figure 1**) as highly selective inhibitor of HDAC3.^41^ Compound **6** contains an *ortho*-methylthioben-zamide Zn^2+^-binding group and exhibits slow-binding mechanism A for HDAC3 inhibition, with *K*_i_ ~24 nM (calculated from kinetic data).^41^ HDACs 1 and 2 are inhibited on endpoint experiments with IC_5_0 values of 20 μM and 31 μM, respectively, but no kinetic analyses are provided for those enzymes. Even though there is a large difference in IC_50_ values based on discontinuous experiments, interrogation of the inhibition of HDACs 1 and 2 in a continuous fashion might be warranted.

HDACs 1–3 form multiprotein complexes in the cell with different sensitivity to inhibitors.^34^ Here, stabilization of the HDAC3/NCoR2 interaction with the “molecular glue” InsP_6_ led to changes in kinetic data that were compound-specific and led to a substantial increase of the measured potency after pre-incubation. These effects might also appear when studying HDACs 1 and 2 in the presence of co-repressors, which would add a new layer of complexity to inhibitor selectivity assessments. Therefore, future HDAC probe characterization may benefit not only from kinetic analysis against the free enzymes, but also against reconstituted complexes. One potential solution may be to measure binding to a library of HDAC complexes by time-resolved Förster resonance energy transfer (TR-FRET), as proposed recently.^37^ These analyses will ensure the development of robust and reproducible tool compounds for HDAC research, which are of importance for the development of epigenetic therapies against neurodegeneration and immune disorders.

In conclusion, we show that entinostat (**3**) is a nanomolar inhibitor of HDACs 1–3 following mechanism B of slow-binding kinetics and that acylhydrazide **5** exhibits subclass differences in kinetic behavior with a preference for inhibition of HDAC3. Importantly, we report that RGFP966 (**4**) is a potent slow-binding inhibitor of HDACs 1–3 and not an HDAC3-selective probe. It is our hope that these findings will assist in future experimental design towards the elucidation of class I HDAC function and subsequent drug development.

## ASSOCIATED CONTENT

### Supporting Information

The Supporting Information is available free of charge on the ACS Publications website.

Supporting figures, schemes and tables; supporting methods, experimental procedures, HPLC traces, and copies of NMR spectra (PDF).

## AUTHOR INFORMATION

### Notes

The authors declare no competing financial interests.

## ACKNOWLEDGMENT

This project has received funding from the European Research Council (ERC) under the European Union’s Horizon 2020 Research and Innovation Programme (grant agreement number CoG-725172–*SIRFUNCT;* C.A.O.).

## ABBREVIATIONS

DAD: deacetylase activation domain
HDAC: histone deacetylase
*k*_obs_: apparent first-order rate constant of equilibration
NCoR2: nuclear receptor co-repressor 2

