## Supporting Information for "Determination of Slow-binding HDAC Inhibitor Potency and Subclass Selectivity"

**Table of contents**

Supporting figures S2

Supporting methods S6

Synthetic procedures S9

Supporting references S11

NMR spectra S12

HPLC traces S15

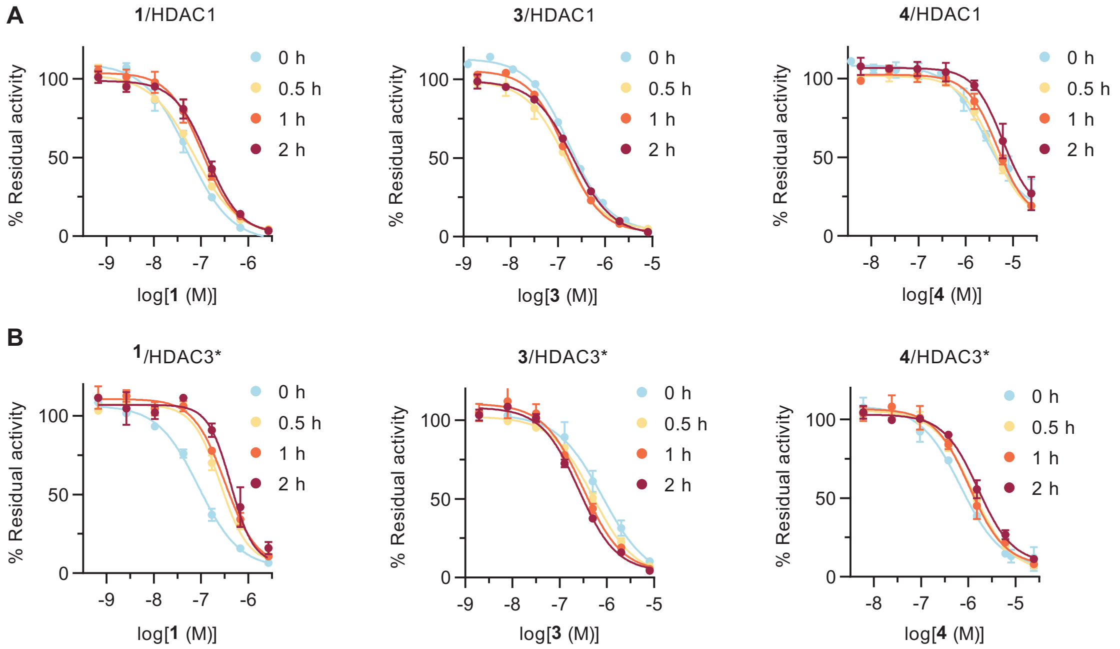


C

| SAHA (**1**) | HDAC1 | HDAC2 | HDAC3* |
|---|---|---|---|
| 0 h: $IC_{50}$ (nM) | 52.3 ± 0.4 | 82 ± 1 | 109 ± 1 |
| 0.5 h: $IC_{50}$ (nM) | 71.8 ± 0.4 | 86.5 ± 0.7 | 259 ± 2‡ |
| 1 h: $IC_{50}$ (nM) | 106 ± 1 | 105 ± 1 | 322 ± 3‡ |
| 2 h: $IC_{50}$ (nM) | 136 ± 1 | 120 ± 2 | 426 ± 5‡ |

| Entinostat (**3**) | HDAC1 | HDAC2 | HDAC3* |
|---|---|---|---|
| 0 h: $IC_{50}$ (nM) | 175 ± 1 | 238 ± 1 | 464 ± 2 |
| 0.5 h: $IC_{50}$ (nM) | 136 ± 1 | 174 ± 1 | 503 ± 3‡ |
| 1 h: $IC_{50}$ (nM) | 145 ± 1 | 108 ± 1 | 311 ± 4‡ |
| 2 h: $IC_{50}$ (nM) | 209 ± 1 | 121 ± 1 | 255 ± 2‡ |

**Figure S1. Discontinuous inhibition of HDACs 1–3 (relative to Figure 2).** (A) HDAC1 inhibition curves with and without pre-incubation. (B) HDAC3 inhibition curves with and without pre-incubation. All were measured in buffer without reducing agents or surfactants. Loss of potency for SAHA (**1**) was attributed to instability of the mixture. (C) Numeric $IC_{50}$ data of compounds **1** and **3**. All data represent mean ± SD, *n* ≥ 2. *HDAC3 incubated with the deacetylase activation domain (DAD) of NCoR2. ‡Data obtained in buffer without reducing agents or surfactants (see **Supporting Figure S2**).

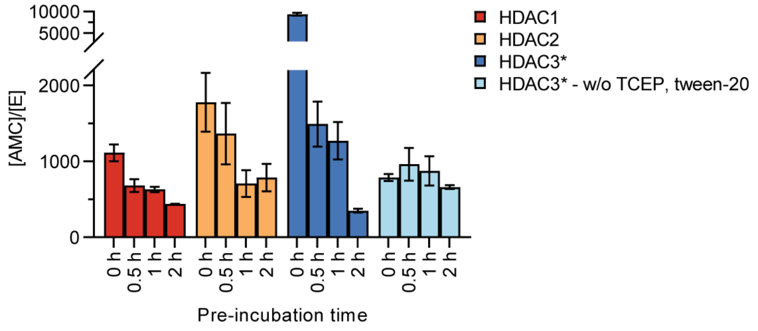


**Figure S2. Enzyme activity after pre-incubation.** Concentration of free AMC measured after pre-incubation and subsequent 30 min reaction, relative to the initial concentration of enzyme employed ([E]). HDAC3 was tested in standard HEPES buffer [50 mM HEPES/Na, 100 mM KCl, 0.001% (v/v) tween-20, 0.2 mM TCEP, pH 7.4, dark blue bars], and in HEPES buffer without TCEP and tween-20 [50 mM HEPES/Na, 100 mM KCl, pH 7.4, light blue bars]. The loss of activity in standard HEPES buffer made it difficult to analyze pre-incubation data, and therefore all HDAC3 pre-incubation experiments were performed in HEPES buffer without TCEP and tween-20. *HDAC3 incubated with the DAD of NCoR2. Data represent mean ± SD, *n* ≥ 2.

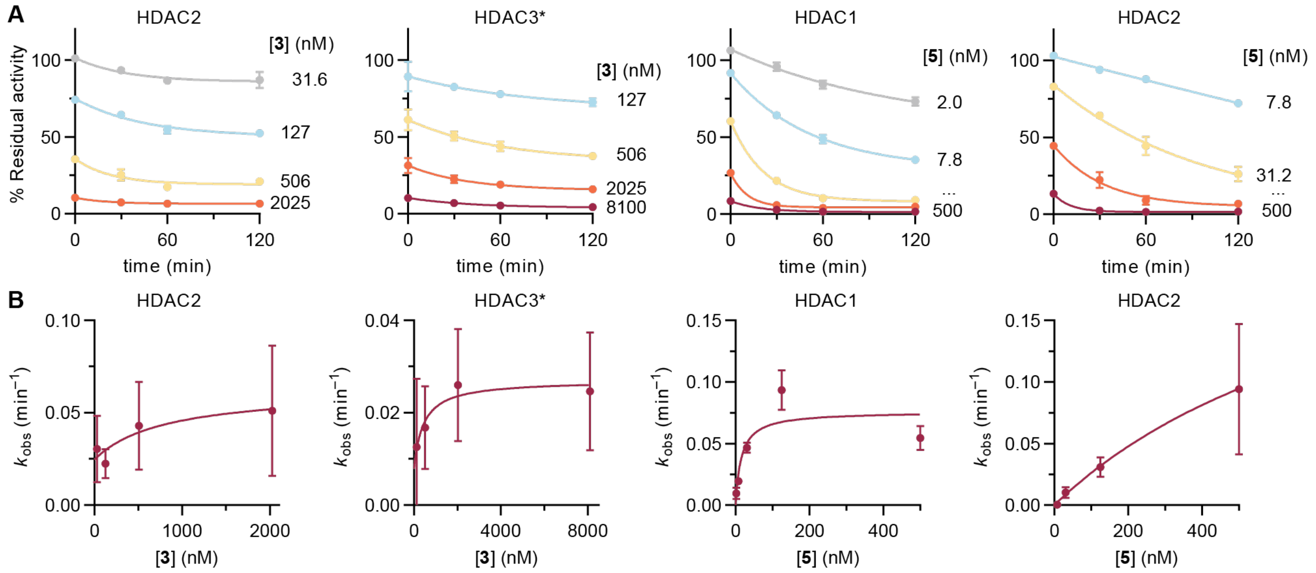


C

| Entinostat (3) | HDAC1 | HDAC2 | HDAC3* |
|---|---|---|---|
| $k_2$ (min$^{-1}$) | N.D. | 0.04 ± 0.03 | 0.019 ± 0.007 |
| $k_{-2}$ (min$^{-1}$) | N.D. | 0.02 ± 0.01 | 0.008 ± 0.008 |
| $K_{i,1}$ (nM) | N.D. | ~190 | ~93 |
| $K_i$ (nM) | N.D. | ~73 | ~27 |
| $t_{½}$ (min) | N.D. | ≥ 28 | ≥ 88 |

| **5** | HDAC1 | HDAC2 | HDAC3* |
|---|---|---|---|
| $k_2$ (min$^{-1}$) | 0.08 ± 0.03 | 0.27 ± 0.08 | N.D. |
| $k_{-2}$ (min$^{-1}$) | ~7 × $10^{-10}$ | ~1 × $10^{-10}$ | N.D. |
| $K_{i,1}$ (nM) | ~4 | 220 ± 100 | N.D. |
| $K_i$ (nM) | –† | –† | N.D. |
| $t_{½}$ (min) | > $10^5$ | > $10^5$ | N.D. |

**Figure S3. Fitting of pre-incubation data to slow-binding inhibitor kinetics.** (A) Data from dose-response inhibition assays of entinostat (**3**) and compound **5** plotted as measured enzyme activity vs. pre-incubation time, and fitted to one-phase exponential decay functions. (B) Secondary plots of $k_{obs}$ vs. concentration of inhibitor, and fitting to mechanism B of slow-binding kinetics. (C) Calculated kinetic parameters relative to mechanism B of slow-binding kinetics. †$K_i$ not determined due to $k_{-2}$ data approaching 0. *HDAC3 incubated with the DAD of NCoR2. N.D.: not determined due to pre-incubation data not fitting exponential decay.

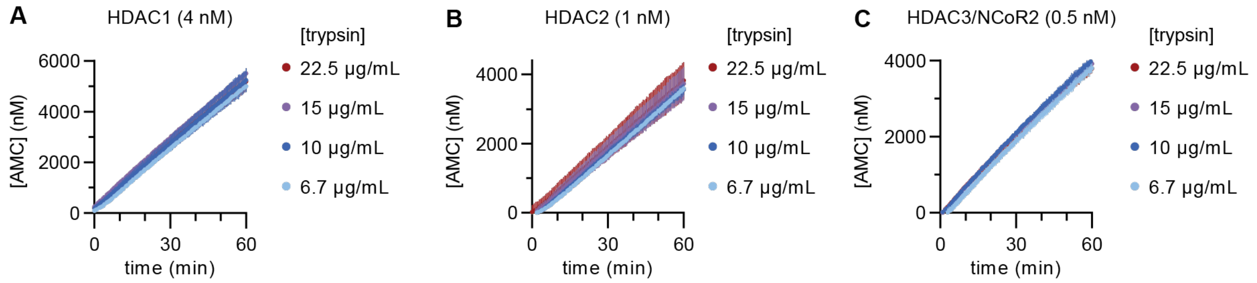


**Figure S4. Optimization of continuous assay conditions.** (A) HDAC1 data set at 4 nM concentration. (B) HDAC2 data set at 1 nM concentration. (C) HDAC3/NCoR2 data set at 0.1 nM concentration. All trypsin concentrations afford similar reaction rates, indicating that HDAC activity is the rate-limiting step of the coupled assay. Data represent mean ± SD of a single experiment with internal duplicate.

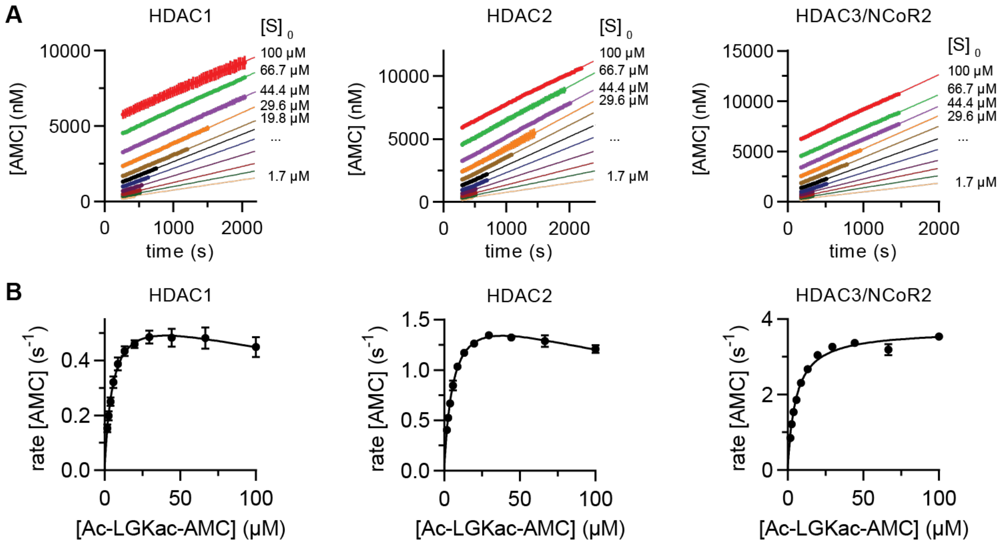


C

Substrate: Ac-LGKac-AMC

| | $K_M$ (µM) | $k_{cat}$ ($s^{-1}$) | $K_i$ (µM) | $k_{cat}/K_M$ ($M^{-1}\cdot s^{-1}$) |
|---|---|---|---|---|
| HDAC1 | 5.4 ± 0.8 | 0.62 ± 0.04 | 300 ± 99 | $(115 \pm 18)\cdot 10^3$ |
| HDAC2 | 6.1 ± 0.5 | 1.76 ± 0.06 | 250 ± 39 | $(291 \pm 24)\cdot 10^3$ |
| HDAC3/NCoR2 | 5.4 ± 0.3 | 3.72 ± 0.06 | – | $(690 \pm 40)\cdot 10^3$ |

**Figure S5. Enzyme kinetic parameters.** (A) Sample progression curves. Data represent mean ± SD of a single experiment with internal duplicate. (B) Initial rate data fitting to the Michaelis-Menten equation (HDAC3/NCoR2) or to the Michaelis-Menten equation with substrate inhibition at high concentrations (HDACs 1 and 2). Rate data are already divided by the initial concentration of enzyme. Data represent mean ± SEM of two independent experiments. (C) Calculated enzyme kinetic parameters. Data represent mean ± SEM of two independent experiments. HDAC3/NCoR2 data are reproduced from a previous study.[1]

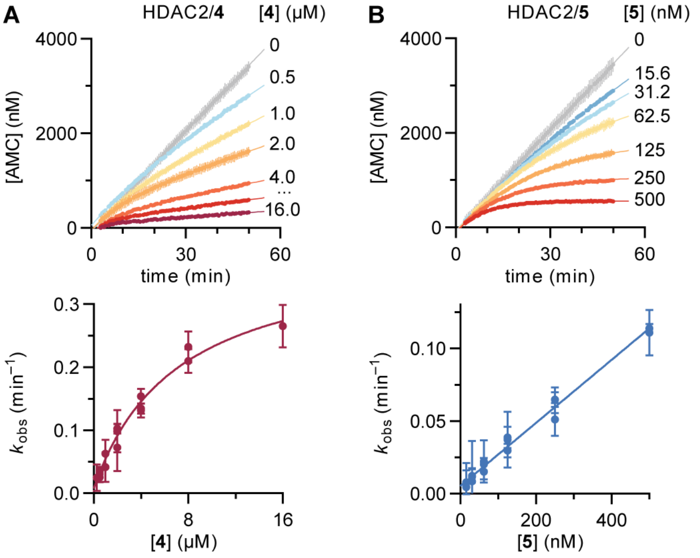


C

Alternative analysis of mechanism A

| **5** | HDAC1 | HDAC2 |
|---|---|---|
| $k_2/K_{i,1}$ ($nM^{-1}\cdot min^{-1}$) | $(1.7 \pm 0.1)\cdot 10^{-3}$ | $(9.3 \pm 0.3)\cdot 10^{-4}$ |
| $k_{-2}$ ($min^{-1}$) | $(3 \pm 3)\cdot 10^{-3}$ | $(5 \pm 2)\cdot 10^{-3}$ |

**Figure S6. Continuous inhibition of HDAC2 by RGFP966 (4) and compound 5.** (A) Continuous assay progression curves (top graph) and $k_{obs}$ secondary plot (bottom graph, mechanism B) relative to compound **4**. (B) Continuous assay progression curves (top graph) and $k_{obs}$ secondary plot (bottom graph, mechanism A) relative to compound **5**. Data represent mean ± SEM of individual experiments, and each experiment was performed at least twice. See **Table 1** for numerical data. (C) Numerical data corresponding to fitting inhibition of HDACs 1 and 2 by compound **5** to **Eq. S13**, where mechanism B is considered but $K_{i,1}$ lays over the [I] range studied ($K_{i,1}$ >> [I]) and cannot be determined.

**A**

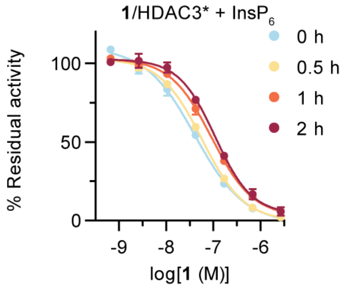


**B**

| SAHA (1) | HDAC3* + InsP$_6$ |
|---|---|
| 0 h: IC$_{50}$ (nM) | 40.8 ± 0.2 |
| 0.5 h: IC$_{50}$ (nM) | 58.1 ± 0.1 |
| 1 h: IC$_{50}$ (nM) | 91.7 ± 0.4 |
| 2 h: IC$_{50}$ (nM) | 109 ± 1 |

| RGFP966 (4) | HDAC3* + InsP$_6$ | Sel. HDAC3† |
|---|---|---|
| 0 h: IC$_{50}$ (μM) | 0.415 ± 0.001 | 4.3 |
| 0.5 h: IC$_{50}$ (μM) | 0.168 ± 0.002 | 18.7 |
| 1 h: IC$_{50}$ (μM) | 0.234 ± 0.002 | 15.7 |
| 2 h: IC$_{50}$ (μM) | 0.291 ± 0.003 | 10.3 |

| Entinostat (3) | HDAC3* + InsP$_6$ |
|---|---|
| 0 h: IC$_{50}$ (nM) | 457 ± 2 |
| 0.5 h: IC$_{50}$ (nM) | 60.6 ± 0.2 |
| 1 h: IC$_{50}$ (nM) | 30.5 ± 0.3 |
| 2 h: IC$_{50}$ (nM) | 14.7 ± 0.1 |

| 5 | HDAC3* + InsP$_6$ | Sel. HDAC3† |
|---|---|---|
| 0 h: IC$_{50}$ (nM) | 3.40 ± 0.01 | 11.7 |
| 0.5 h: IC$_{50}$ (nM) | 1.02 ± 0.01 | 6.5 |
| 1 h: IC$_{50}$ (nM) | 0.95 ± 0.02 | 3.5 |
| 2 h: IC$_{50}$ (nM) | 0.95 ± 0.02 | 1.5 |

**C**

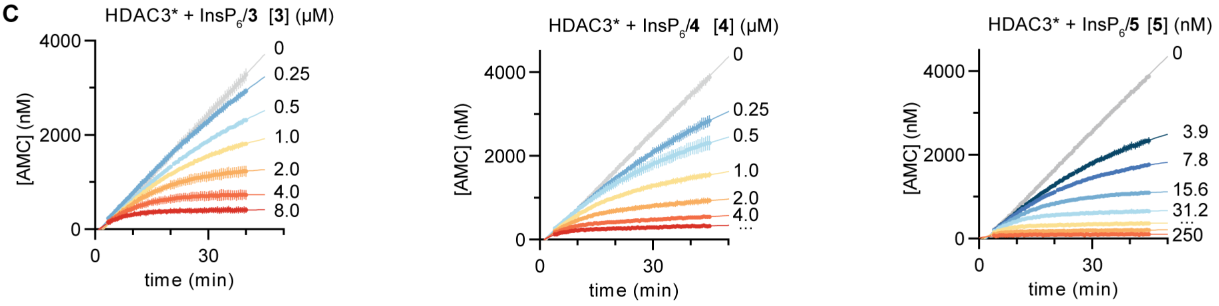


**D**

| HDAC3* + InsP$_6$ | Entinostat (3) | RGFP966 (4) | 5 |
|---|---|---|---|
| $k_2$ (min$^{-1}$) | 0.44 ± 0.08 | 0.57 ± 0.07 | 0.54 ± 0.06 |
| $k_{-2}$ (min$^{-1}$) | ~4 × 10$^{-3}$ | ~4 × 10$^{-10}$ | ~4 × 10$^{-10}$ |
| $K_{i,1}$ (nM) | 2400 ± 700 | 1600 ± 400 | 17 ± 5 |
| $K_i$ (nM) | ~24 | –† | –† |
| t$_{½}$ (min) | ≥ 160 | > 10$^5$ | > 10$^5$ |

**Figure S7. Inhibition of HDAC3/NCoR2 in the presence of InsP$_6$.** (A) Curves relative to SAHA (**1**) after pre-incubation with enzyme and InsP$_6$ for 0, 0.5, 1, or 2 h, followed by addition of substrate and 30 min reaction. Data represent mean ± SD, *n* = 2. (B) IC$_{50}$ data and selectivity of HDAC3 inhibition. †Selectivity calculated vs. the second most inhibited enzyme (**4**: HDAC2, **5**: HDAC1), by transforming IC$_{50}$ data into $K_i$ and calculating the ratio (see data analysis in the **Supporting Methods**). (C) Continuous assay progression curves relative to compounds **3**, **4**, and **5**. (D) Calculated kinetic constants ($k_n$), inhibitor constants ($K_n$), and dissociation half-lives (t$_{½}$) of compounds **3**, **4**, and **5** in the presence of InsP$_6$, corresponding to mechanism B of slow-binding kinetics. All experiments were carried out in buffer without reducing agents or surfactants (see **Supporting Figure S2** and **Supporting Methods**). *HDAC3 incubated with the deacetylase activation domain (DAD) of NCoR2.

### Supporting Methods

#### Assay materials

HDAC assays were performed in black low-binding 96-well microtiter plates (Corning half-area wells, Fischer Scientific, cat. # 3686), with duplicate series in each assay and each assay performed at least twice. Control wells without enzyme were included in each plate. All experiments were performed in HEPES buffer [50 mM HEPES/Na, 100 mM KCl, 0.001% (v/v) tween-20, 0.2 mM tris(2-carboxyethyl)phosphine (TCEP), pH 7.4][2] with 0.5 mg/mL bovine serum albumin (BSA, Sigma-Aldrich, cat. #: A7030) unless otherwise indicated. Phytic acid sodium salt hydrate ($InsP_6$, Sigma-Aldrich, cat. #: 68388) and trypsin (Sigma-Aldrich, cat. #: T1426) were of commercial source. The following recombinant enzymes were acquired from BPS Bioscience (San Diego, CA): HDAC1 (full length, C-terminal His tag, cat. #: 50051, lot. #: 200108-A), HDAC2 (full length, C-terminal FLAG tag, cat. #: 50052, lot. #: 140324-1), HDAC3/NCoR2 (full length, C-terminal His tag, with DAD domain of NCor2, N-terminal GST tag, cat. #: 50003, lot. #: 190327). The following inhibitors were of commercial source: SAHA (**1**, Sigma-Aldrich, cat. #: SML0061), entinostat (**3**, MS-275, Sigma-Aldrich, cat. #: EPS002), and RGFP966 (**4**, Sigma-Aldrich, cat. #: 16917), and the purity of corresponding stocks was verified by HPLC during the course of the current study (see **HPLC traces**). Stocks were prepared in DMSO (10–40 mM), concentration of inhibitor **5** was determined by NMR by addition of an internal maleic acid standard, and concentration of peptide substrates was determined based on absorbance [$\varepsilon_{326}$(Ac-Lys-AMC) = 17783 $M^{-1}\cdot cm^{-1}$][3] using a Thermo Scientific NanoDrop$^{C}$ instrument. Assay concentration of substrates and inhibitors were obtained by dilution from DMSO stock solutions in buffer, and the appropriate concentration of enzyme was obtained by dilution of the stock provided by the supplier. $InsP_6$ stocks were prepared in fresh in assay buffer (62 mg/mL ≈ 50 mM). Fluorescence recordings were performed in a FLUOstar Omega plate reader (BMG Labtech). Data analysis was performed using GraphPad Prism 9.

#### Dose-response end-point inhibition assays, no pre-incubation

Inhibitors (72.9 μM–0.048 nM or 900–0.048 nM, 3-fold dilutions; **1**: 2.7 μM–0.66 nM, 4-fold dilutions) were incubated in HEPES buffer with substrate (Ac-Leu-Gly-Lys(Ac)-AMC, 20 μM) and enzyme (HDAC1: 4 nM, HDAC2: 3 nM, HDAC3/NCoR2: 1 nM) for 30 min at 37 ºC in a volume of 25 μL/well. For experiments with $InsP_6$, inhibitors (**1**: 2.7 μM–0.66 nM, **2**: 8.1 μM–2.0 nM, **3**: 24.3 μM–5.9 nM, or **4**: 500–0.12 nM, 4-fold dilutions) were incubated in HEPES buffer without tween-20 or TCEP (50 mM HEPES/Na, 100 mM KCl, pH 7.4) with substrate (Ac-Leu-Gly-Lys(Ac)-AMC, 20 μM), $InsP_6$ (100 μM) and enzyme (HDAC3/NCoR2: 1 nM) for 30 min at 37 ºC in a volume of 25 μL/well. Thereafter, a trypsin solution (0.4 mg/mL in corresponding HEPES buffer) was added (25 μL/well, for a final volume of 50 μL/well), the plate was left standing for 15 min at room temperature before fluorescence recording. Background fluorescence from wells without enzyme was subtracted from all data, and the residual activity (%) was calculated relative to control wells without inhibitor. All experiments were performed at least twice and with internal replicate wells. Data were fitted to sigmoidal functions with variable slope (**Eq. S1**, 4 parameters, *h*: Hill slope) in order to calculate $IC_{50}$ values, and the Cheng-Prusoff equation (**Eq. S2**) was employed to transform $IC_{50}$ data into inhibitory constants ($K_i$) for selectivity calculations, where $K_M$ values were as determined (see **Supporting Fig. S5C**). Selectivity ratios were determined by dividing the corresponding $K_i$ values.

$$\mathrm{Res.\,activity} = \mathrm{Res.\,activity_{bottom}} + \frac{\mathrm{Res.\,activity_{top}} - \mathrm{Res.\,activity_{bottom}}}{1 + 10^{(\log \mathrm{IC_{50}} - \log[\mathrm{I}])h}} \quad (\mathbf{Eq.\,S1})$$

$$K_\mathrm{i} = \frac{\mathrm{IC_{50}}}{1 + \frac{[\mathrm{S}]}{K_\mathrm{M}}} \quad (\mathbf{Eq.\,S2})$$

**Dose-response end-point inhibition assays, with pre-incubation**
Inhibitors (**1**: 2.7 μM–0.66 nM, **2**: 8.1 μM–2.0 nM, **3**: 24.3 μM–5.9 nM, or **4**: 500–0.12 nM, 4-fold dilutions) were incubated for 0.5, 1, or 2 h at 37 ºC in HEPES buffer with enzyme (HDAC1: 5 nM for 0.5 h, 10–12 nM for 1 h, 13–15 nM for 2 h; or HDAC2: 4 nM for 0.5 h, 9 nM for 1 h, 10–12 nM for 2 h), followed by addition of substrate (Ac-Leu-Gly-Lys(Ac)-AMC, 20 μM) for a volume of 25 μL/well (pre-incubation performed in 20 μL, and concentrations reported for the 25 μL reaction volume with substrate). Due to instability of the enzyme preparation, assays with HDAC3/NCoR2 were performed in HEPES buffer without tween-20 or TCEP (50 mM HEPES/Na, 100 mM KCl, pH 7.4), and with 5 nM enzyme concentration for all pre-incubation times (see **Supporting Fig. S2**). Control dose-response experiments were performed without pre-incubation to ensure inhibitor $IC_{50}$ data did not change with the modification of the buffer. For experiments with $InsP_6$, inhibitors were incubated in HEPES buffer without tween-20 or TCEP with $InsP_6$ (100 μM) and enzyme (HDAC3/NCoR2: 1 nM), followed by addition of substrate (Ac-Leu-Gly-Lys(Ac)-AMC, 20 μM). Reactions were then incubated for 30 min at 37 ºC, then a trypsin solution (0.4 mg/mL in the corresponding HEPES buffer) was added (25 μL/well, for a final volume of 50 μL/well), and the plate was left standing for 15 min at room temperature before fluorescence recording. All experiments were performed at least twice and with internal replicate wells. Data was analyzed as described for end-point experiments without pre-incubation.
In order to determine kinetic parameters from pre-incubation data, residual enzyme activity (%) was fitted to **Eq. S3** of one-phase exponential decay, and the calculated $k_{obs}$ parameters were analyzed as for continuous HDAC inhibition assays (see below).

$$\text{Res. activity} = (\text{Res. activity}_0 - \text{Res. activity}_\infty)e^{(-k_{obs}t)} + \text{Res. activity}_\infty \quad (\textbf{Eq. S3})$$

**Determination of enzyme kinetic parameters**
HDAC1 (4 nM), HDAC2 (2 nM), or HDAC3 (1 nM) were incubated with substrate (Ac-Leu-Gly-Lys(Ac)-AMC, 100–1.73 μM, 1.5-fold dilutions) and trypsin (10 μg/mL) at room temperature in HEPES buffer at a final volume of 50 μL/well. Fluorescence was read every 30 s for 60 min, and assay progression curves (up to 10% substrate conversion) were fitted to linear regressions to provide reaction rates ($\nu_0$). All experiments were performed at least twice and with internal replicate wells. A standard curve of free AMC fluorescence was measured under the same conditions and used to transform fluorescence units into AMC concentration. HDAC3 data were fitted to the Michaelis-Menten equation (**Eq. S4**), where $[E]_0$ is the initial concentration of HDAC and [S] is the initial concentration of substrate, which provided the $K_M$ and $k_{cat}$ kinetic parameters (see **Supporting Fig. S5A–C**).

$$\frac{\nu_0}{[E]_0} = \frac{k_{cat}[S]}{K_M + [S]} \quad (\textbf{Eq. S4})$$

Data corresponding to HDACs 1 and 2 were fitted to the Michaelis-Menten equation with substrate inhibition at high concentrations (**Eq. S5**), where $K_i$ is the inhibitory constant of the substrate (see **Supporting Fig. S5C**).

$$\frac{\nu_0}{[E]_0} = \frac{k_{cat}[S]}{K_M + [S]\left(1 + \frac{[S]}{K_i}\right)} \quad (\textbf{Eq. S5})$$

**Optimization of continuous assay conditions**
HDAC1 (4 nM, 2 nM, or 1 nM), HDAC2 (4 nM, 2 nM, or 1 nM), or HDAC3/NCoR2 (2 nM, 1 nM, or 0.5 nM) were incubated with substrate (Ac-Leu-Gly-Lys(Ac)-AMC, 20 μM) and trypsin (22.5–6.7 μg/mL, 1.5-fold dilutions) at room temperature in HEPES buffer at a final volume of 50 μL/well. Fluorescence was read every 30 s, and assay curves were plotted to determine reaction rates (see **Supporting Fig. S4**).

**Continuous HDAC inhibition assays**

For a final volume of 50 µL, substrate (Ac-Leu-Gly-Lys(Ac)-AMC, 20 µM), trypsin (6.7 µg/mL), inhibitor (16000–125 nM or 500–3.9 nM, 2-fold dilutions) and enzyme (HDAC1: 4 nM, HDAC2: 1 nM, HDAC3: 0.5 nM) were added in HEPES buffer to a microtiter plate and immediately placed in the plate reader. For experiments with $InsP_6$, inhibitors, substrate (Ac-Leu-Gly-Lys(Ac)-AMC, 20 µM), $InsP_6$ (100 µM) and enzyme (HDAC3/NCoR2: 0.5 nM) were added in HEPES buffer without tween-20 or TCEP (50 mM HEPES/Na, 100 mM KCl, pH 7.4). *In situ* fluorophore release was monitored by fluorescence readings every 30 s for 60 min at 25 °C, and fluorescence data was transformed into concentration of free fluorophore ([AMC], nM) using a standard curve recorded under the same conditions. All experiments were performed at least twice and with internal replicate wells. Data was then fitted to **Eq. S6** in order to determine the apparent first-order rate constant for establishment of the enzyme-inhibitor equilibrium ($k_{obs}$). In this equation, $v_{in}$ and $v_{ss}$ are the initial and final steady-state velocities of each experiment, respectively.

$$[\mathrm{AMC}] = v_{ss}t + \frac{v_{in} - v_{ss}}{k_{obs}}\left(1 - \mathrm{e}^{-k_{obs}t}\right) \quad (\mathbf{Eq.\,S6})$$

Assuming competitive inhibition (Cheng-Prusoff model), $k_{obs}$ and inhibitor concentration were fitted to either mechanism A (linear relationship, **Eq. S7**) or mechanism B (hyperbolic relationship, **Eq. S8**) of slow kinetics to afford the corresponding kinetic constants. Here, the magnitude of $K_{i,1}$ determines the mechanism, since mechanism A is the limiting case where $K_{i,1}$ >> [I] and only a single step of binding can be determined from the data fitting. $K_M$ values were as determined (see **Supporting Fig. S5C**).

$$k_{obs} = k_1\left(1 + \frac{[\mathrm{S}]}{K_\mathrm{M}}\right)[\mathrm{I}] + k_{-1} \quad (\mathbf{Eq.\,S7})$$

$$k_{obs} = \frac{k_2}{[\mathrm{I}] + K_{i,1}\left(1 + \frac{[\mathrm{S}]}{K_\mathrm{M}}\right)}[\mathrm{I}] + k_{-2} \quad (\mathbf{Eq.\,S8})$$

Whenever possible, inhibitor constants ($K_i$) and enzyme-inhibitor dissociation half-life ($t_{½}$) values were estimated using either **Eq. S9** and **Eq. S10** (mechanism A) or **Eq. S11** and **Eq. S12** (mechanism B), respectively.[4] Since $k_{-1}$ is not determined here for mechanism B inhibitors, only the lower half-life limit is calculated, where $k_{-1}$ >> $k_{-2}$,$k_2$, and $t_{½}$ ~ $\ln(2)/k_{-2}$.

$$K_i = \frac{k_{-1}}{k_1} \quad (\mathbf{Eq.\,S9})$$

$$t_{½} = \frac{\ln(2)}{k_{-1}} \quad (\mathbf{Eq.\,S10})$$

$$K_i = K_{i,1}\frac{k_{-2}}{k_2 + k_{-2}} \quad (\mathbf{Eq.\,S11})$$

$$t_{½} = \frac{\ln(2)}{\frac{k_{-1} \cdot k_{-2}}{(k_{-1} + k_2 + k_{-2})}} \quad (\mathbf{Eq.\,S12})$$

Data relative to mechanism A (compound **5** vs. HDAC1,2) was also analyzed according to a more generic mechanism B for which $K_{i,1}$ cannot be determined within the concentration range of inhibitor tested ($K_{i,1}$ >> [I]). In that case, [I] is simplified from the denominator of **Eq. S8**, which affords **Eq. S13**.

$$k_{\mathrm{obs}} = \frac{k_2}{K_{\mathrm{i,1}}\left(1+\frac{[\mathrm{S}]}{K_{\mathrm{M}}}\right)}[\mathrm{I}] + k_{-2} \quad (\mathbf{Eq.\,S13})$$

Fitting $k_{obs}$ data to **Eq. S13** provides mechanism B data included in **Supporting Figure S6C**.

### Synthetic Procedures

#### General synthetic procedures

All reagents and solvents were of analytical grade, and they were used without further purification as obtained from commercial suppliers. Reactions were monitored by thin-layer chromatography (TLC), using silica gel-coated plates (analytical $SiO_2$-60, with F-254), and visualized under UV light and/or with a $KMnO_4$ solution. Alternatively, reactions were followed by high performance liquid chromatography-mass spectrometry (HPLC-MS) on a Waters Acquity UPLC system with a C18 Phenomenex Kinetex column (1.7 µm, 50 × 2.1 mm, 100 Å) and photodiode array (PDA) and single quadrupole (SQ) detectors, using a linear gradient of eluent I (0.1% HCOOH in $H_2O$) and eluent II (0.1% HCOOH in MeCN) rising linearly from 0% to 95% of II during t = 0.20–4.80 min at a flow rate of 0.6 mL/min. Column chromatography was performed using a Buchi PureFlash system with EcoFlex silica cartridges. Preparative reversed-phase HPLC purification was performed on a Agilent 1260 LC system equipped with a C18 Phenomenex Luna column (5 µm, 250 × 20 mm, 100 Å), and diode array and evaporative light scattering (ELS) detectors. Linear gradients of eluent III ($H_2O$/MeCN/TFA, 95:5:0.1) and eluent IV (0.1% TFA in MeCN) $t$ = 5–35 min were employed for the purification, with a flow rate of 20 mL/min. The purity of collected fractions and of the final compound were then checked on an Agilent 1260 Infinity II series UPLC system with a C18 Infinity Poroshell 120 column [100 × 3.0 mm, 2.7 µm] and a diode array detector, using a 0–50% linear gradient of eluent IV in eluent III during t = 1–11 min with a flow rate of 1.2 mL/min at 40 °C. Nuclear magnetic resonance (NMR) spectra were recorded at 298 K on a Bruker Avance III HD equipped with a cryogenically cooled probe ($^1$H NMR and $^{13}$C NMR recorded at 600 and 151 MHz, respectively). Chemical shifts are reported in ppm relative to deuterated solvent as internal standard. Assignments of NMR spectra are based on 2D correlation spectroscopy (COSY, HSQC and HMBC spectra).

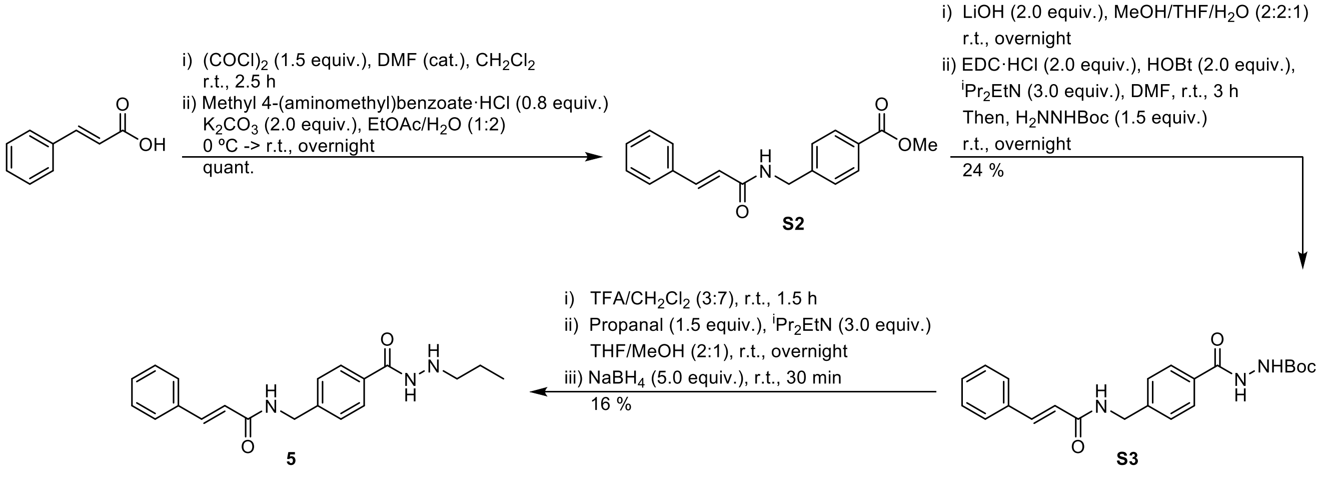


**Scheme S1. Synthesis of compound 5.**

**Methyl 4-(cinnamamidomethyl)benzoate (S2).** *trans*-Cinnamic acid (2.00 g, 13.5 mmol, 1.3 equiv.) was dissolved in anh. $CH_2Cl_2$ (170 mL) under $N_2$ atmophere. To this solution, oxalyl chloride (1.7 mL, 20.3 mmol, 2.0 equiv.) and DMF (3 drops) were added dropwise via syringe. The mixture was stirred at room temperature for 2.5 h, condensed under vacuum, brought to 0 °C and diluted with an ice-cold EtOAc/$H_2O$ mixture (1:2, 150 mL). Then, $K_2CO_3$ (3.73 g, 27.0 mmol, 2.6 equiv.) and methyl 4-(aminomethyl)benzoate·HCl (2.09 g, 10.4 mmol, 1.0 equiv.) were added portion-wise, and the reaction was allowed to reach room temperature and stir overnight. The aqueous and organic phases were separated, the aqueous phase was extracted with EtOAc (2 × 100 mL), and the combined organic phases were washed with aq. HCl (0.1 M, 100 mL), sat. aq. $NaHCO_3$ (100 mL), and brine (100 mL). The organic phase was dried over $MgSO_4$, filtered, and concentrated under vacuum to afford the title compound as off-white solid (3.44 g), which was used in the next step without further purification.

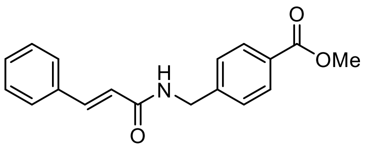


$^1$H NMR (600 MHz, DMSO-d6): δ 8.72 (t, $J$ = 6.0 Hz, 1H, NH), 7.96 – 7.89 (m, 2H, $C_{Ar}\underline{H}C_{Ar}COO$), 7.60 – 7.55 (m, 2H, $C_{Ar}\underline{H}C_{Ar}CH$), 7.49 (d, $J$ = 15.8 Hz, 1H, $C_{Ar}CH$), 7.45 – 7.34 (m, 5H, $C_{Ar}\underline{H}C_{Ar}HC_{Ar}CH$, $C_{Ar}\underline{H}C_{Ar}HC_{Ar}HC_{Ar}CH$, $CH_2C_{Ar}C_{Ar}\underline{H}$), 6.71 (d, $J$ = 15.8 Hz, 1H, $C\underline{H}CONH$), 4.49 (d, $J$ = 6.0 Hz, 2H, $CH_2$), 3.84 (s, 3H, $CH_3$).

$^{13}$C NMR (151 MHz, DMSO-d6): δ 166.1 (CONH), 165.1 (COO), 145.1 ($CH_2\underline{C_{Ar}}$), 139.2 ($C_{Ar}\underline{C}H$), 134.8 ($\underline{C_{Ar}}CH$), 129.5 ($\underline{C_{Ar}}COO$), 129.3 ($C_{Ar}H$), 128.9 ($C_{Ar}H$), 128.2 ($C_{Ar}H$), 127.6 ($C_{Ar}H$), 127.4 ($C_{Ar}H$), 121.80 ($\underline{C}HCONH$), 52.0 ($CH_3$), 42.0 ($CH_2$).

LCMS: $m/z$ 296.09 ($[M + H]^+$, calcd. for $C_{18}H_{18}NO_3^+$: 296.13).

***tert*-Butyl 2-(4-(cinnamamidomethyl)benzoyl)hydrazine-1-carboxylate (S3).** Compound **S2** (1.0 g, 3.39 mmol, 1.0 equiv.) was dissolved in a MeOH/THF/$H_2O$ mixture (2:2:1, 25 mL), and LiOH (162 mg, 6.78 mmol, 2.0 equiv.) was added portion-wise. The reaction was allowed to stir overnight, after which the reaction mixture was concentrated in vacuo, diluted with $CH_2Cl_2$/MeOH (9:1, 200 mL), washed with aq. HCl (1 M, 100 mL) and brine (100 mL). The combined aqueous layer was extracted with $CH_2Cl_2$/MeOH (9:1, 2 × 50 mL), and the combined organic layer was dried over $MgSO_4$, filtered, and concentrated under vacuum to afford 4-(cinnamamidomethyl)benzoic acid as white solid (894 mg, 94%). The obtained solid (894 mg, 3.18 mmol, 1.0 equiv.) was dissolved in DMF (34 mL) under $N_2$ atmosphere, followed by addition of 1-hydroxybenzotriazole (HOBt, 859 mg, 6.36 mmol, 2.0 equiv.), $^iPr_2EtN$ (1.66 mL, 9.54 mmol, 3.0 equiv.), and *N*-(3-dimethylaminopropyl)-*N′*-ethylcarbodiimide hydrochloride (EDC·HCl, 1.22 g, 6.36 mmol, 2.0 equiv.). The activation was allowed to proceed for 3 h at room temperature, followed by addition of *tert*-butyl carbazate (630 mg, 4.77 mmol, 1.5 equiv.) and reaction overnight at room temperature. Then, the mixture was concentrated in vacuo, diluted with EtOAc (200 mL), and washed with aq. HCl (0.1 M, 2 × 100 mL), sat. aq. $NaHCO_3$ (100 mL), and brine (100 mL). The organic layer was dried over $MgSO_4$, filtered, concentrated under vacuum, and purified by automated flash column chromatography (0–40% EtOAc in heptane) to afford the title compound as white solid (321 mg, 24% from **S2**).

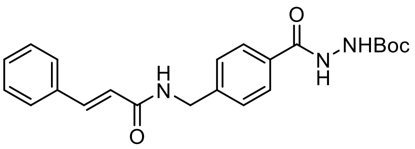


$^1$H NMR (600 MHz, DMSO-d6) δ 10.15 (s, 1H, $C_{Ar}CONH$), 8.88 (s, 1H, NHCOO), 8.69 (t, $J$ = 6.0 Hz, 1H, $N\underline{H}CH_2$), 7.83 (d, $J$ = 7.9 Hz, 2H, $C_{Ar}\underline{H}C_{Ar}CONH$), 7.60 – 7.55 (m, 2H, $C_{Ar}\underline{H}C_{Ar}CH$), 7.49 (d, $J$ = 15.8 Hz, 1H, $C_{Ar}CH$), 7.45 – 7.35 (m, 5H, $C_{Ar}\underline{H}C_{Ar}HC_{Ar}CH$, $C_{Ar}\underline{H}C_{Ar}HC_{Ar}HC_{Ar}CH$, $CH_2C_{Ar}C_{Ar}\underline{H}$), 6.71 (d, $J$ = 15.8 Hz, 1H, $C\underline{H}CONH$), 4.47 (d, $J$ = 6.0 Hz, 2H, $CH_2$), 1.43 (s, 9H, $C(CH_3)_3$).

$^{13}$C NMR (151 MHz, DMSO-d6) δ 165.8 (CONHNH), 165.1 ($CH\underline{C}ONH$), 155.5 (NHCOO), 143.3 ($CH_2\underline{C_{Ar}}$), 139.1 ($C_{Ar}\underline{C}H$), 134.8 ($\underline{C_{Ar}}CH$), 131.1 ($\underline{C_{Ar}}CONH$), 129.5 ($C_{Ar}H$), 128.9 ($C_{Ar}H$), 127.5 ($C_{Ar}H$),

127.4 ($C_{Ar}H$), 127.2 ($C_{Ar}H$), 127.1 ($C_{Ar}H$), 121.9 ($\underline{C}HCONH$), 79.1 ($\underline{C}(CH_3)_3$), 42.0 ($CH_2$), 28.1 ($C(\underline{C}H_3)_3$).
LCMS: *m/z* 396.30 ($[M + H]^+$, calcd. for $C_{22}H_{26}N_3O_4^+$: 396.19).

***N*-(4-(2-Propylhydrazine-1-carbonyl)benzyl)cinnamamide (5).** Compound **S3** (32 mg, 0.081 mmol, 1.0 equiv.) was stirred in $CH_2Cl_2/CF_3COOH$ (7:3, 3.3 mL) for 1.5 h, after which volatiles were removed under a $N_2$ stream. The resulting crude oil mixture was dispersed in heptane/toluene (1:1) and concentrated under vacuo twice to afford a white solid. The obtained solid was dissolved in THF/MeOH (2:1, 3 mL) under $N_2$ atmosphere, followed by addition of $^iPr_2EtN$ (84 µL, 0.48 mmol, 6.0 equiv.) and propionaldehyde (15 µL, 0.21 mmol, 2.6 equiv.). The reaction mixture was stirred overnight, then placed in an ice bath followed by addition of $NaBH_4$ (31 mg, 0.81 mmol, 10.0 equiv.). The mixture was stirred for additional 30 min at room temperature, quenched with $H_2O$, concentrated in vacuo, and purified by preparative reverse phase HPLC (0–40% eluent IV in eluent III) to afford the title compound as white powder after lyophilization (4.5 mg, 97% purity, 16% yield).

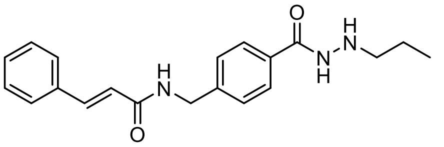

$^1H$ NMR (600 MHz, DMSO-d6) δ 10.91 (s, 1H, $C_{Ar}CONH$), 8.72 (t, *J* = 6.0 Hz, 1H, $CON\underline{H}CH_2$), 7.86 – 7.80 (m, 2H, $C_{Ar}\underline{H}C_{Ar}CONH$), 7.60 – 7.55 (m, 2H, $C_{Ar}\underline{H}C_{Ar}CH$), 7.48 (d, *J* = 15.8 Hz, 1H, $C_{Ar}CH$), 7.45 – 7.35 (m, 5H, $C_{Ar}\underline{H}C_{Ar}HC_{Ar}CH$, $C_{Ar}\underline{H}C_{Ar}HC_{Ar}HC_{Ar}CH$, $CH_2C_{Ar}C_{Ar}\underline{H}$), 6.71 (d, *J* = 15.8 Hz, 1H, $C\underline{H}CONH$), 4.47 (d, *J* = 6.0 Hz, 2H, $CONHC\underline{H_2}$), 2.98 (t, *J* = 7.5 Hz, 2H, $NHNHC\underline{H_2}$), 1.58 (h, *J* = 7.4 Hz, 2H, $C\underline{H_2}CH_3$), 0.93 (t, *J* = 7.4 Hz, 3H, $CH_2C\underline{H_3}$).
$^{13}C$ NMR (151 MHz, DMSO-d6) δ 165.3 (CONHNH), 165.1 ($CH\underline{C}ONH$), 144.0 ($CH_2\underline{C_{Ar}}$), 139.1 ($C_{Ar}\underline{C}H$), 134.8 ($\underline{C_{Ar}}CH$), 130.1 ($\underline{C_{Ar}}CONH$), 129.5 ($C_{Ar}H$), 128.9 ($C_{Ar}H$), 127.6 ($C_{Ar}H$), 127.5 ($C_{Ar}H$), 127.3 ($C_{Ar}H$), 121.9 ($\underline{C}HCONH$), 52.1 ($NHNHCH_2$), 42.0 ($CONH\underline{C}H_2$), 19.0 ($\underline{C}H_2CH_3$), 11.2 ($CH_2\underline{C}H_3$).
LCMS: *m/z* 338.18 ($[M + H]^+$, calcd. for $C_{20}H_{24}N_3O_2^+$: 338.19).

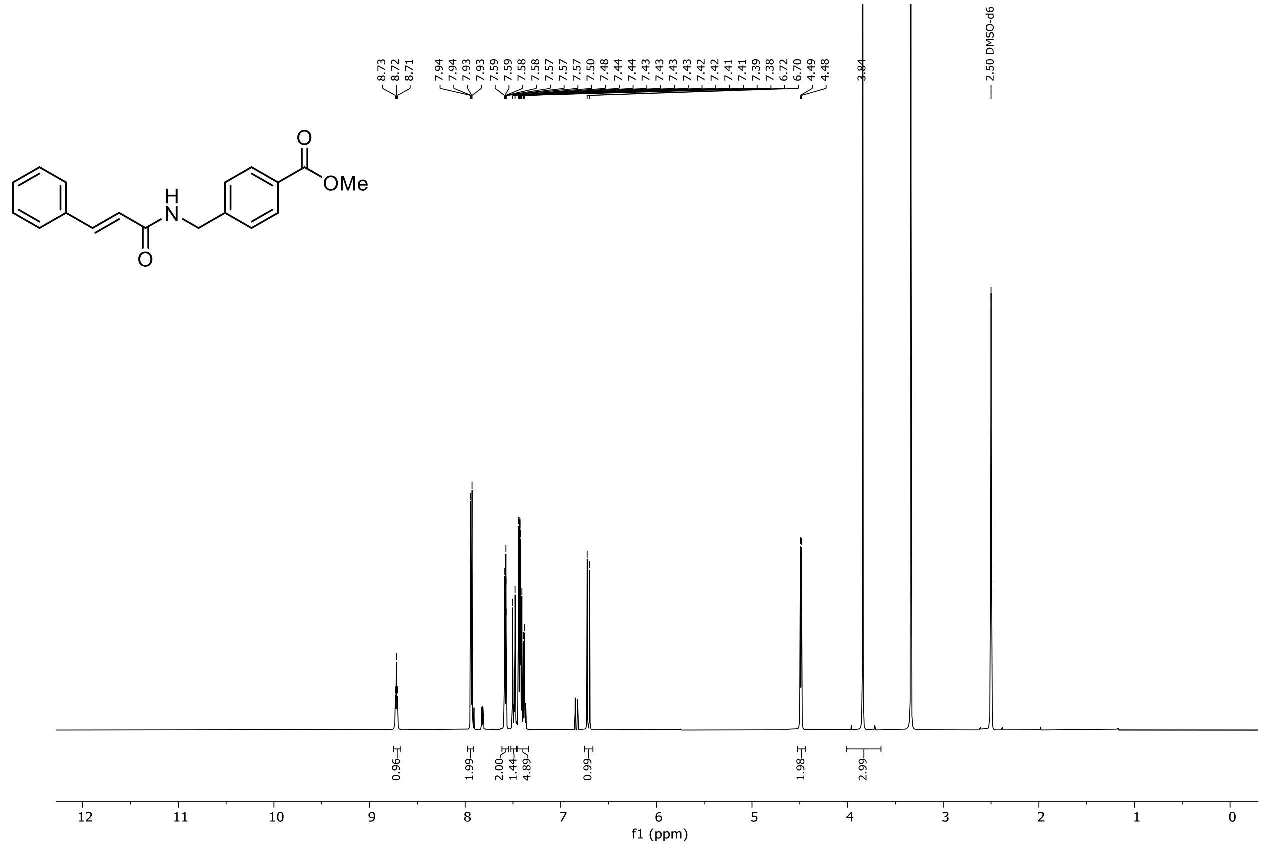


$^{1}H$ NMR spectrum of compound **S2**.

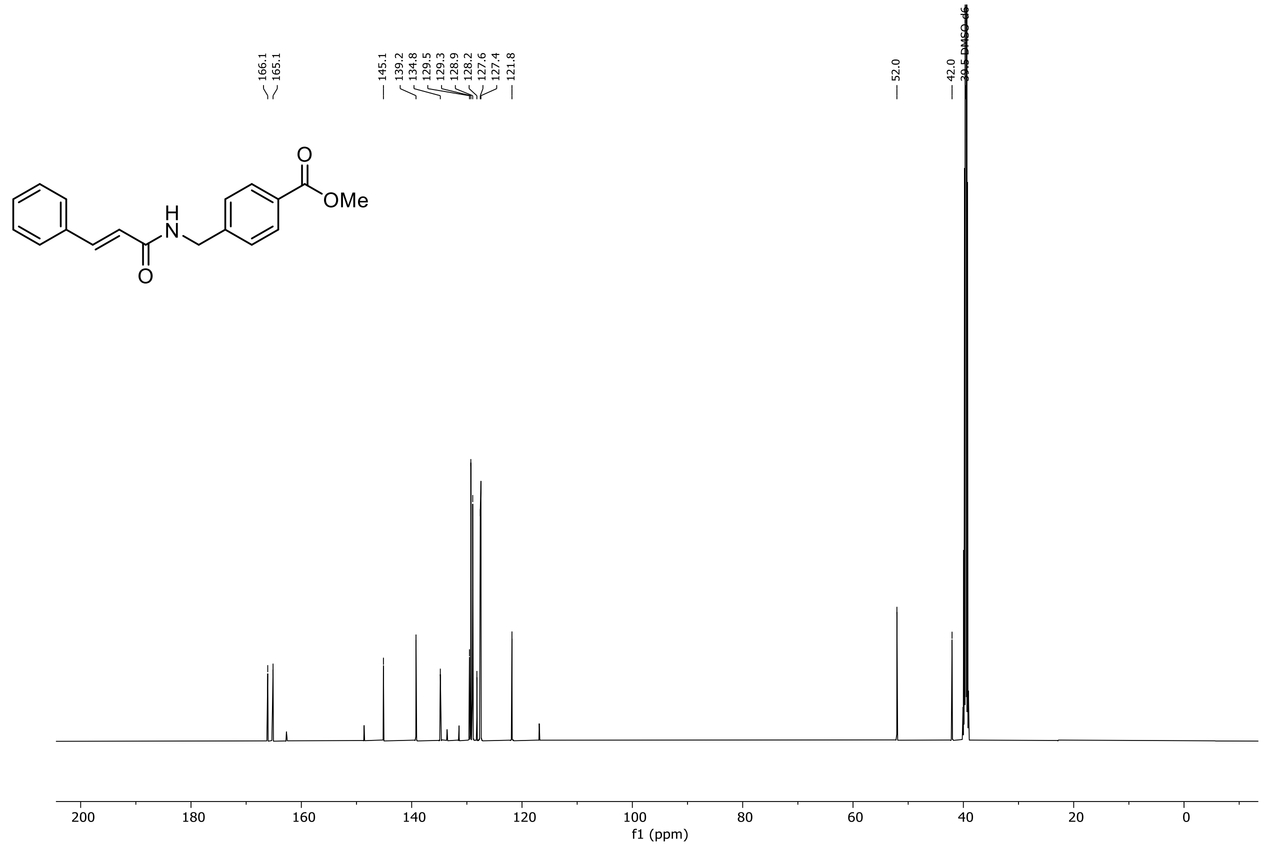


$^{13}C$ NMR spectrum of compound **S2**.

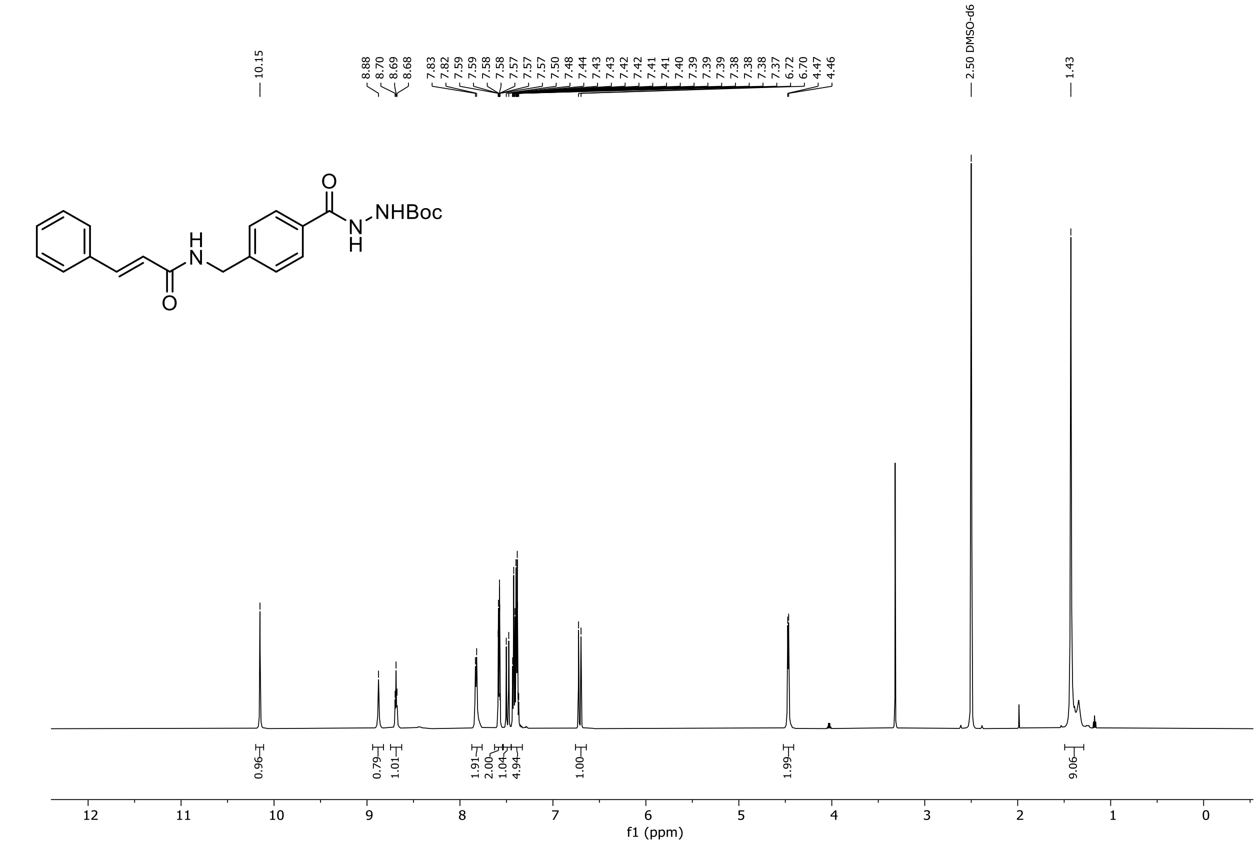


[1]H NMR spectrum of compound **S3**.

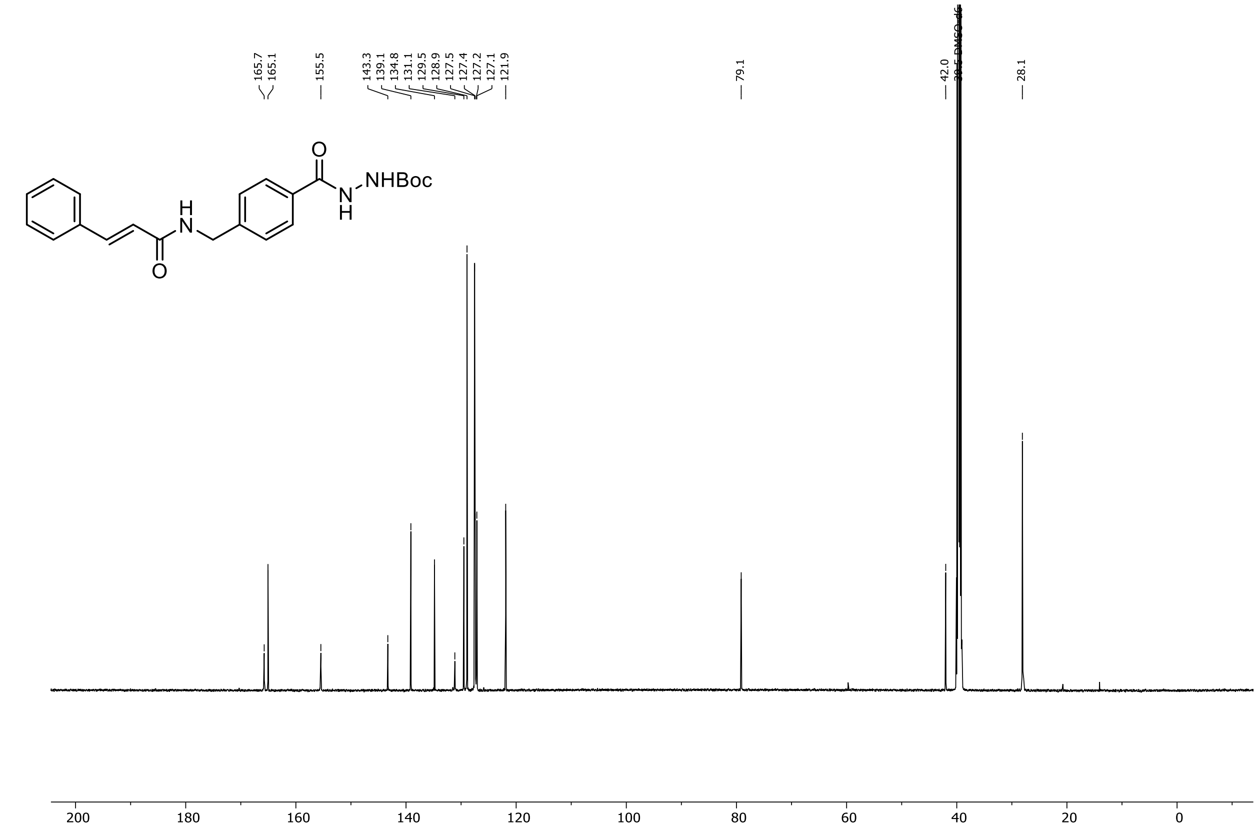


[13]C NMR spectrum of compound **S3**.

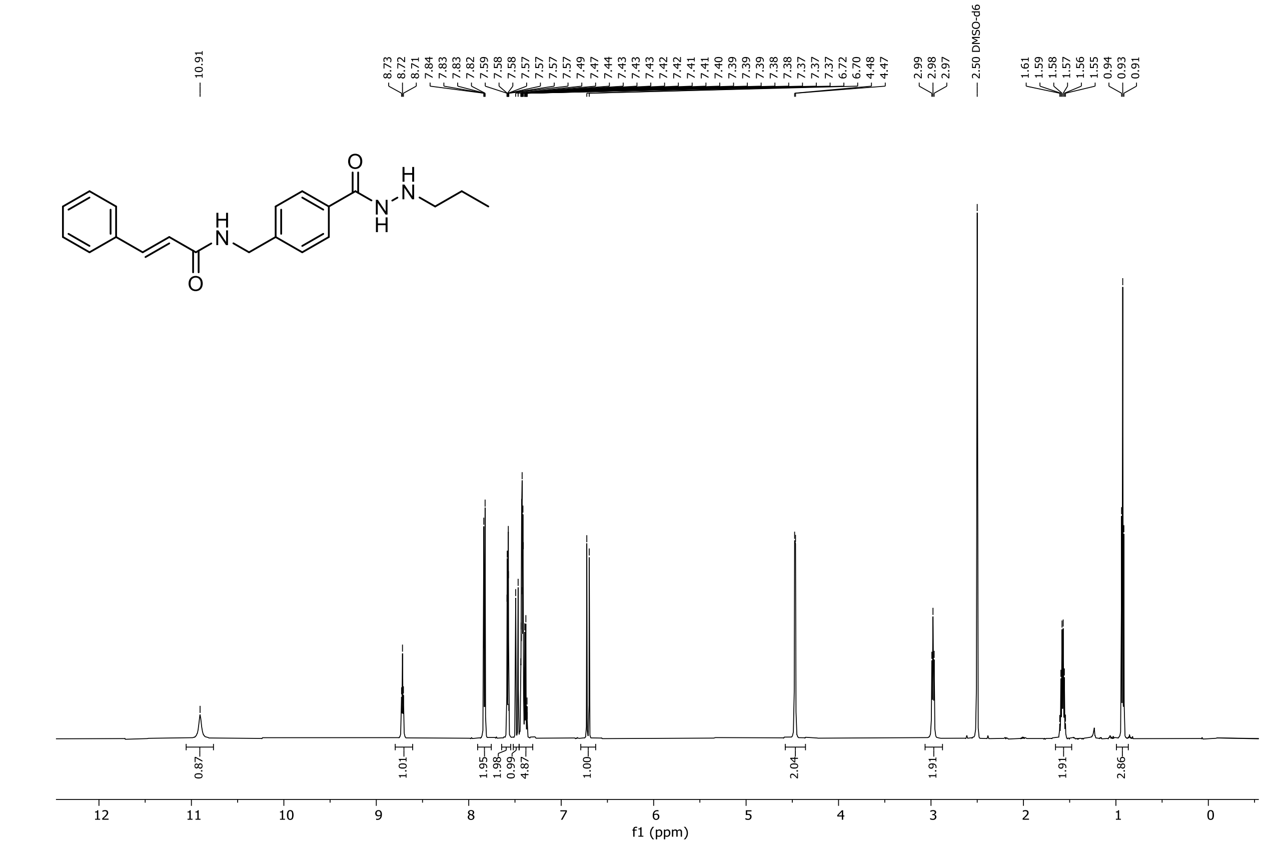


$^{1}$H NMR spectrum of compound **5**.

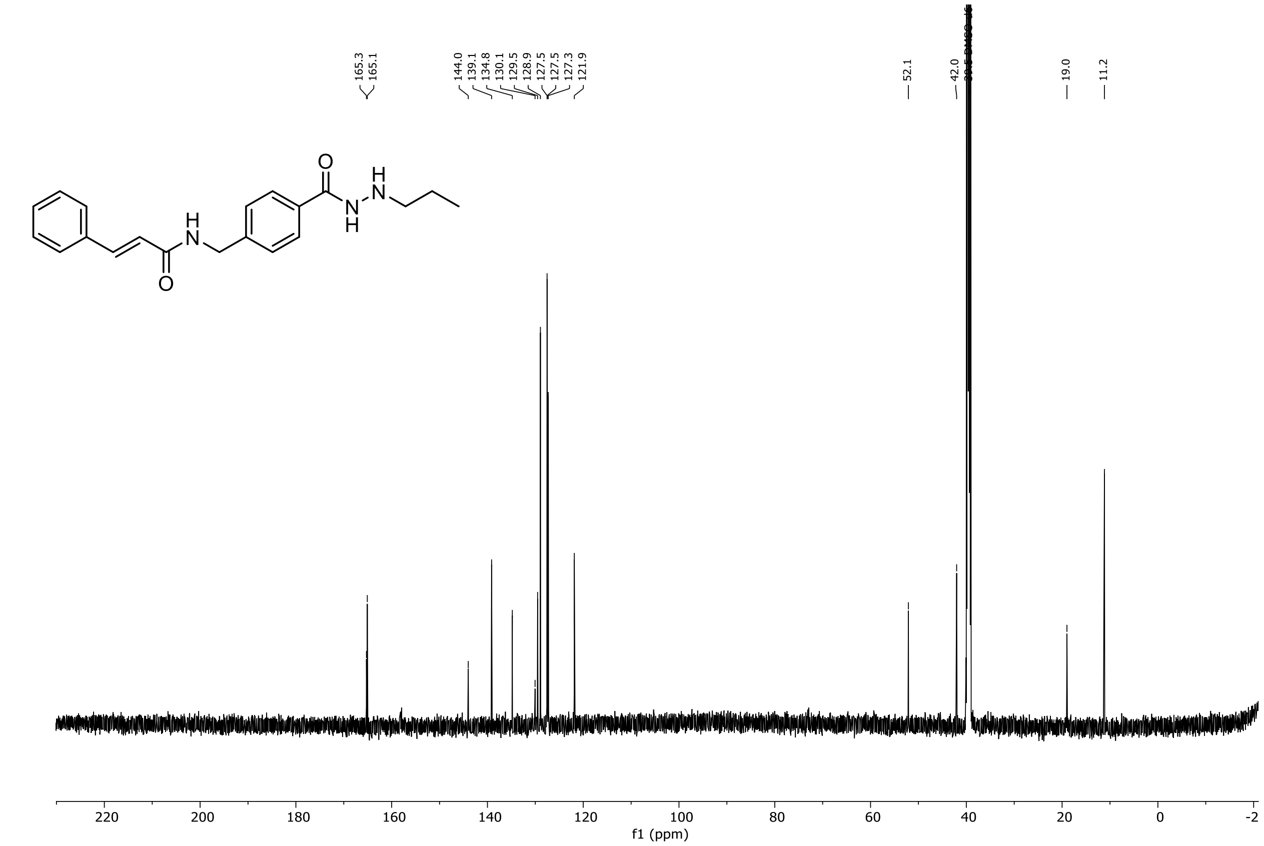


$^{13}$C NMR spectrum of compound **5**.

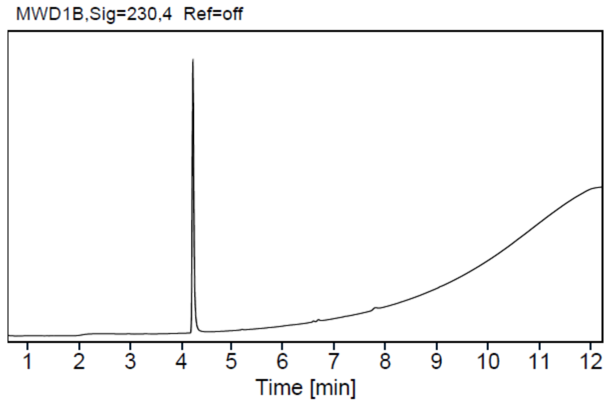


HPLC purity trace of compound **1** (0–95% gradient).

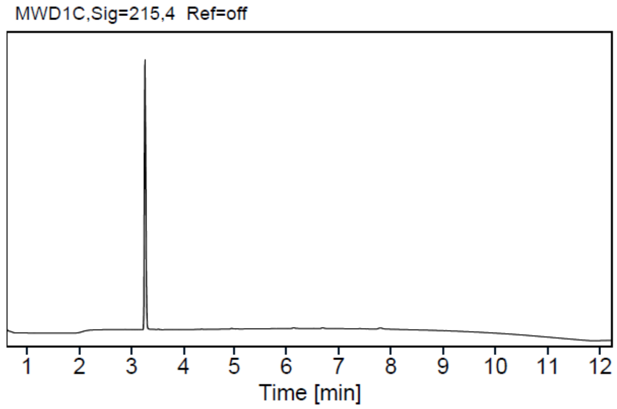


HPLC purity trace of compound **3** (0–95% gradient).

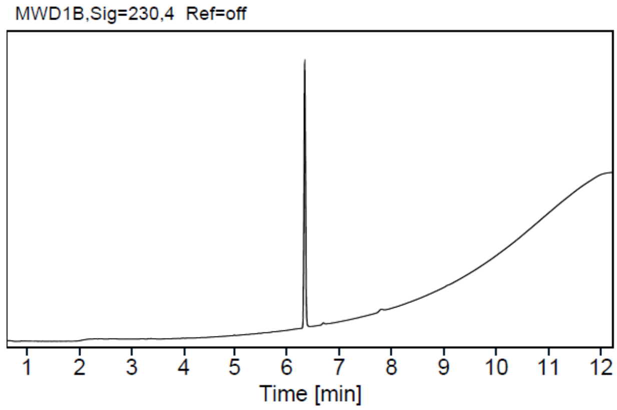


HPLC purity trace of compound **4** (0–95% gradient).

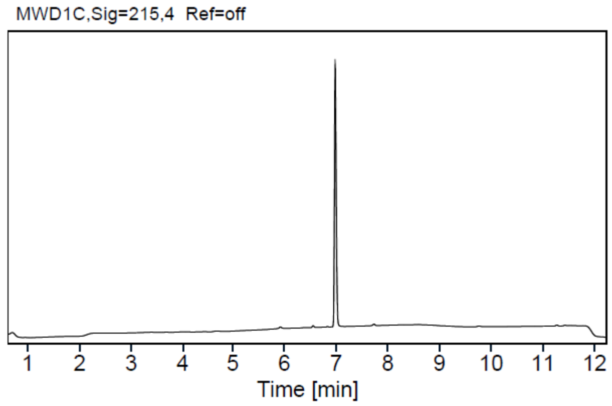


HPLC purity trace of compound **5** (0–50% gradient).
